# A quantitative census of millions of postsynaptic structures in a large electron microscopy volume of mouse visual cortex

**DOI:** 10.64898/2026.02.19.706834

**Authors:** Benjamin D. Pedigo, Bethanny P. Danskin, Rachael Swanstrom, Erika Neace, Sven Dorkenwald, Nuno Maçarico da Costa, Casey M. Schneider-Mizell, Forrest Collman

## Abstract

Neurons display remarkable sub-cellular specificity in their synaptic targeting, which varies by cell type—for example, excitatory neurons prefer to target the spines of other excitatory cells. Modern electron microscopy connectomes enable the study of this sub-cellular specificity and its context in a circuit at unprecedented scale and resolution. However, this scale has also made it challenging to create accurate and efficient methods for classifying and segmenting fine cell components (including spines) across entire volumes. Here, we present a cost-efficient computational pipeline for classifying postsynaptic targets and segmenting structures such as spines. Our method relies only on having a mesh representation of a neuron and avoids processing image or segmentation data directly. Instead, we leverage tools from geometry processing to create features capturing the local geometry of a neuron’s surface. We couple this core technique with computational and storage optimizations, enabling reliable deployment over hundreds of thousands of neurons for a few hundred dollars in cloud compute cost. We then show that a simple classifier trained on the MICrONS mouse visual cortex dataset can use these mesh-based features to accurately classify synapses as targeting somas, dendritic shafts, or spines (weighted F1 score 0.961). Using this pipeline, we create a map of the postsynaptic structures at over 207.3 million synapses in MICrONS. We present an overview of this census of postsynaptic targeting, finding expected patterns (e.g., excitatory neurons preferentially targeting excitatory spines) as well as unexpected exceptions (e.g., Layer 5 near-projecting and Layer 6 corticothalamic cells often connecting to excitatory neuron shafts). We also demonstrate that these tools can be used to detect spines receiving multiple synaptic inputs, revealing surprising variability in their frequency across cells even within a cell type. We make our postsynaptic target predictions available for study, as well as the code for the computational pipeline and cloud deployment. Beyond MICrONS, we find that the model generalizes well to the H01 connectome without retraining (weighted F1 score 0.949), indicating that these tools will be useful in future connectomics reconstructions. More generally, our work demonstrates that representations derived from neuronal meshes can be a scalable and generalizable primitive for describing morphologies.

## 1 Introduction

### 1.1 Spines and subcellular targeting

For as long as neuroanatomy has been a field, dendritic spines have been appreciated as a salient feature of neural morphology. Spines are small protrusions from dendrites found on many neuronal types in mammalian nervous systems, and in cortex are the canonical site of excitatory-to-excitatory connections. The functional role of spines has been debated, but they are thought to be a mechanism for biochemical isolation of signals and a method for reaching a broader sample of presynaptic targets [1]. Spines vary greatly in morphology [2], are dynamic and change in response to experience [3, 4], and can change in their density, shape, and dynamics with disease [5].

Many of the previous studies on spine morphologies have leveraged either light microscopy techniques [2, 6] (which typically enables the study of spines across entire arbors of sparsely sampled cells) or electron microscopy in small to moderate sized volumes [7, 8] (which typically enables highly detailed dense reconstructions of a small number of spines). Recent technological developments have enabled a massive scaling up of our ability to collect and analyze large volumetric electron microscopy (EM) data, including in cortex [9–11]. These datasets present the opportunity to examine synapses and their specific target compartments in great detail and in the context of a local circuit. Understanding subcellular targeting patterns in these large reconstructions is also a useful substrate for many downstream analyses. For instance, spine morphology has been shown to be a helpful feature for classifying cell types [12–14], compartment targeting patterns can be used to infer the excitatory or inhibitory nature of presynaptic axon fragments [8, 15], and spine sizes have been used as estimates of synaptic weight [16, 17]. However, with these large reconstructions also come new challenges in analyzing morphology at this scale while still retaining the level of detail required to characterize spines and other postsynaptic structures.

### 1.2 Computational approaches to postsynaptic structure characterization

Many computational approaches have been developed for detecting and/or segmenting spines [13, 14, 18–23]. These tools range in their approach, from heuristics based on skeletonizations and mesh geometry [14, 19] to deep learning pipelines which operate on imagery and/or segmentation [18, 22, 23]. The variety of this work shows that many approaches can be effective at segmenting spines. However, applying such techniques at scale across massive EM datasets is often computationally burdensome and expensive. Some also rely upon having access to auxiliary segmentations of subcellular components such as mitochondria, which may not be available for all datasets [18]. To our knowledge, few of these pipelines have been applied broadly across a dataset at the scale of MICrONS (though see [14]).

Thus, despite this progress, there remained a need for reliable, economical methods for densely segmenting spines at the scale of a cubic-millimeter EM volume. Much of the computational cost in many of the approaches outlined above comes from downloading and processing huge amounts of dense imagery and segmentation data. However, many EM processing pipelines already include tools for creating meshed representations of segmented objects. Meshes describe the surface of objects with a discrete triangulation, which is typically a more compact descriptor of an object’s external shape. This makes meshes an attractive object for more concisely representing and learning on neuron morphologies (some of the aforementioned previous works have also leveraged point or surface representations; see for example [18, 20, 21]).

One added difficulty in working with meshes is that they are irregular domains (unlike images, they have no regular grid structure), so specialized methods are required to generate or learn representations of these objects. Due to their ubiquity in many scientific and computing domains, meshes have received a great deal of attention from the computer graphics and computational geometry literature, and many authors have explored the idea of constructing shape representations on the basis of mesh structure alone. Much of the early work in this space has focused on descriptions based on the eigenstructure of a surface’s Laplace-Beltrami operator (hereafter referred to as the Laplacian), which is a generalization of the Laplace operator to other domains such as meshes. The eigenvectors of the Laplacian can be thought of as a generalization of the Fourier series to arbitrary graphs, and resemble a series of periodic patterns of increasing frequency when visualized on meshes.

Here, we leverage one of these methods known as the heat kernel signature (HKS) [24]. We show that the mesh features generated by this method can be used to create an accurate classifier for postsynaptic shapes. We describe computational modifications to this basic algorithm that allow it to run on millimeter scale EM datasets at low cost. Then, we apply this pipeline to create a census of postsynaptic structures in the MICrONS dataset. This allows us to explore patterns of sub-cellular targeting both within and across cell types in this local circuit. We also develop an extension of these techniques which allows us to find spines which receive multiple synaptic inputs, and again present a census of their occurrence in MICrONS. We expect that the synaptic annotations presented here will be a useful resource for future studies of connectivity in MICrONS, and also show that this technique will enable similar analyses in other large scale neuroanatomy datasets.

## 2 Results

### 2.1 Heat kernel signatures on neuron morphologies

We developed a computational pipeline to predict postsynaptic structure (spine, dendritic shaft, or soma) based purely on the geometry of neuronal meshes, which are ubiquitous in volumetric EM pipelines. Our approach leverages the heat kernel signature (HKS) of Sun *et al*. [24], a tool from computational geometry which generates descriptive features of a mesh’s surface. Conceptually, the HKS feature vector for a mesh vertex is constructed by initializing a unit of heat at that vertex, and tracking how much heat is left at that vertex for each of a set of prespecified timesteps (Figure 1A-B). These heat retention values are concatenated into the HKS vector for that vertex, and rescaled to more evenly weight features at different timescales (Figure 1B). Sun *et al*. [24] showed that these features characterize the local geometry of a surface in a way that allows for matching of points with nearly congruent neighborhoods. Further, these features are intrinsic, meaning they are invariant to rotations, reflections, and translations, which endows models based on the HKS with the same properties without the need for extensive data augmentations.

**Figure 1.**
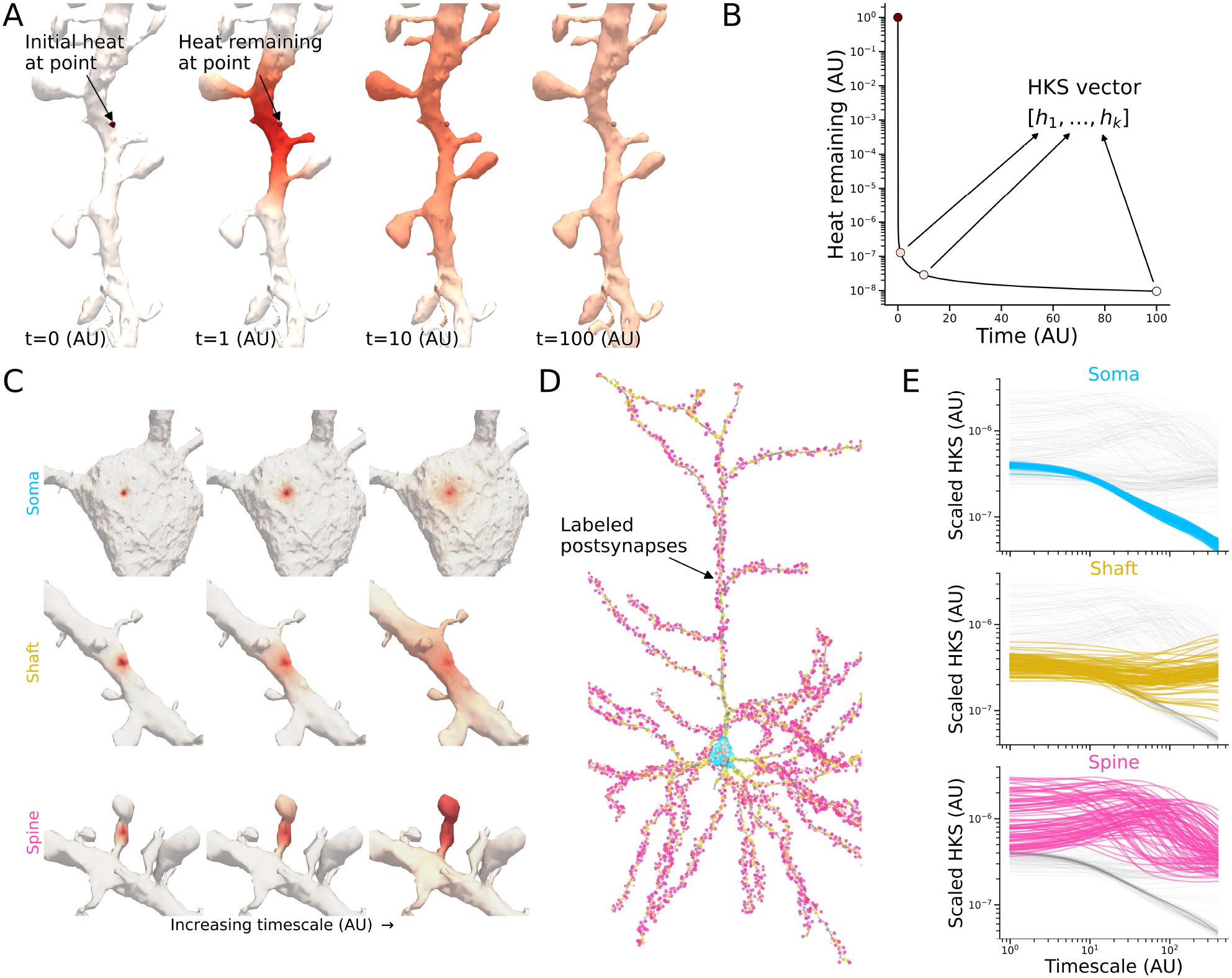
Diagram of heat kernel signature (HKS) computation and its use on neuron surface meshes. A) Conceptually, the HKS is computed by placing a unit of heat at a point on a mesh, and allowing it to diffuse on the mesh surface according to the heat equation. B) Plot of heat remaining at the example point in A as time (AU) evolves. The HKS for a given mesh vertex are the amount of heat remaining at each of a set of timescales, which are assembled into a vector representing that point on the mesh. C) Example heat diffusion on the mesh shown for a postsynaptic site on the soma (top), a dendritic shaft (middle), and a spine (bottom). Note the differential behavior of heat diffusion over time based on the local geometry of the mesh. D) Manual labeling applied to all putative synaptic inputs on an example neuron. E) HKS features for mesh points associated with each of the synapse categories. HKS features have been scaled by the mean value for that timescale for ease of visualization. Note that the scaled HKS features show clear separability in terms of the soma, shaft, and spine categories.

To understand whether these features would be sufficient to characterize postsynaptic structures, we simulated the heat diffusion process and corresponding HKS construction for example points on a pyramidal neuron mesh from the MICrONS dataset [9] (Figure 1C). For these points, heat dissipates slowly from dendritic spines due to their isolated nature, less slowly from the dendritic shaft, and most quickly from the point on the soma. To see whether this pattern held generally across the neuron, we manually labeled all of its input synapses (and their corresponding mesh vertices) as being onto the soma, shaft, or spine (Figure 1D). We found that the HKS features for these vertices clearly separated according to the manual labels (Figure 1E).

### 2.2 Scaling heat kernel signature computation

This motivated us to develop a computational pipeline for computing HKS features at scale in large neuroanatomy volumes such as MICrONS, as well as a classifier for predicting postsynaptic shape from these HKS features. Rather than simulating heat diffusion from each vertex on each mesh (of which there can be millions), the standard scheme for computing heat kernel signatures performs an eigendecomposition on the mesh’s discrete Laplace-Beltrami operator (Laplacian) [24], which is analogous to the well-known graph Laplacian commonly used in machine learning and other fields. This approach solves for the heat trace at multiple timescales and across the entire mesh in one pass rather than repeating a simulation for many points and timescales, and can be truncated to a specified number of eigenpairs to save computation. However, computing even this partial eigendecomposition to the resolution required to resolve spines can be prohibitively expensive for an entire neuron mesh.

We also sought to reduce the storage footprint of the computed features. The HKS yields a feature vector for each of the millions of vertices in a neuron’s mesh, which is costly to store across large datasets. Retaining these HKS features (rather than just the classifications alone) enables rapid classifier iteration and reuse for other tasks (for instance, segmenting axonal boutons; see Figure S1 and section 3) without repeating the computation of feature generation.

Our approach to computing the HKS robustly and at scale leverages several techniques (diagrammed in Figure 2A), including:

- mesh simplification [25] to reduce the number of vertices to compute over,
- overlapping mesh chunks to construct smaller problems to solve while minimizing edge effects,
- use of the robust Laplacian [26] to model heat diffusion while minimizing the effects of imperfections in meshes,
- a band-by-band approach [27] to accelerate the eigendecomposition, and
- mesh agglomeration to locally approximate the heat kernel signatures of similar mesh vertices to reduce storage size.

**Figure 2.**
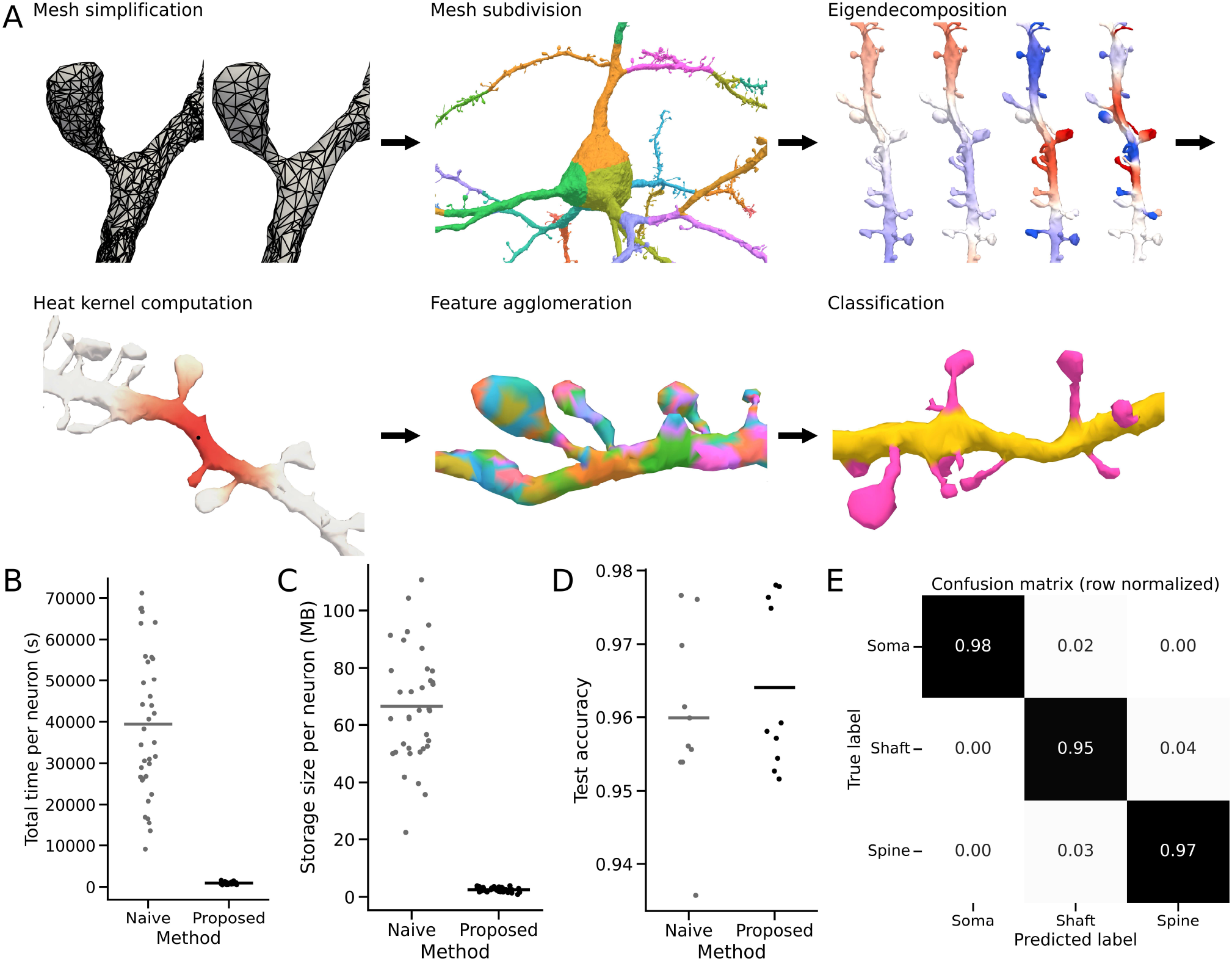
A scalable, accurate pipeline for postsynaptic structure prediction from neuronal mesh data. A) Diagram of key steps in the proposed cloud-deployable pipeline for computing heat kernel signatures on neuronal meshes. Meshes are first simplified, and then broken into overlapping subregions to speed computation (each mesh subdomain can be processed independently). For each subdomain, we compute the eigendecomposition of the mesh’s robust Laplacian operator (example eigenvectors plotted on the mesh surface are shown). These eigenvector/eigenvalue pairs are used to compute the heat kernel signature for all mesh vertices simultaneously. These features are compressed for storage via a connectivity-constrained clustering on the surface of the mesh based on the HKS features, and using the mean feature vector within each cluster as representative of that group of vertices. Classification can then be run on the surface of the mesh using the condensed features. B) Total time to compute the HKS representation for a naive and proposed version of the HKS computation. C) Storage size for saving the HKS embedding for a naive implementation which stores an HKS vector for each mesh vertex, versus our proposed implementation which uses agglomeration as a lossy compression scheme. Both storage sizes are shown after gzip compression. D) Cross-validation accuracy scored on held out neurons with and without the proposed computational modifications to the HKS computation. E) Mean confusion matrix on held out neurons over 5 cross-validation folds.

We describe these approaches in more detail in subsection 4.3.

These modifications to the naive eigendecomposition-based HKS computation saved time and storage space for the HKS vectors, while still enabling us to train an accurate classifier for postsynaptic structure. We profiled the computation time for 36 example neurons with and without our proposed scheme. On average, computation time was reduced 42-fold (Figure 2B). For the same neurons, we also profiled the compressed size of the file storing the HKS features with and without our vector agglomeration scheme, which can be viewed as a form of topology-aware lossy compression. Our approach reduced storage size for these example neurons by 27-fold on average.

Importantly, these improvements came without sacrificing the fidelity required to construct an accurate classifier. We collected manual labels for 110,946 synapses in the MICrONS dataset, and then mapped these synaptic labels to their corresponding mesh vertices and their HKS feature vectors (see Figure S2 for an analysis of the dependence of this classifier on training samples). Using this training data and labels, we trained a random forest classifier [28] to predict postsynaptic structure label from the HKS features. We did not observe a difference in accuracy between the naive and our proposed schemes on held-out neurons, with both having 96.8% average accuracy on held-out neurons (Figure 2D-E).

In summary, we created a highly efficient pipeline for HKS computation from neuronal meshes^1^, a containerized cloud deployment of this pipeline for horizontal scaling using Kubernetes^2^, and a classifier for predicting postsynaptic structure from HKS features ^3^. These tools are all available as open source software on GitHub, enabling their easy application to other datasets. We have also created a web-based demo of these tools to enable researchers to quickly try the proposed techniques on their own data^4^.

### 2.3 Postsynaptic structure prediction across MICrONS

This pipeline allowed us to scale computation of heat kernel signatures and the corresponding postsynaptic target classifications to the MICrONS dataset. We selected 74,728 segmented objects in the dataset that had been previously identified as neurons using a classifier based on perisomatic and nuclear features [12]. We ran our proposed pipeline for HKS computation and postsynaptic structure prediction on these neurons, labeling 207.3 million input synapses in total. Of these, 123.3 million were predicted to be onto spines, 78.7 million were predicted to be onto dendritic shafts, and 5.3 million were predicted to be onto somas. We estimate that the cost of running this pipeline in the commercial cloud on these neurons was less than $500, making it extremely affordable relative to the scale of the dataset. Example neuron prediction maps are shown for various cell types as defined in Elabbady *et al*. [12] in Figure 3. While the model was trained using sparse points (center locations of synapses), the model performs well at segmenting the entire surface of the mesh.

**Figure 3.**
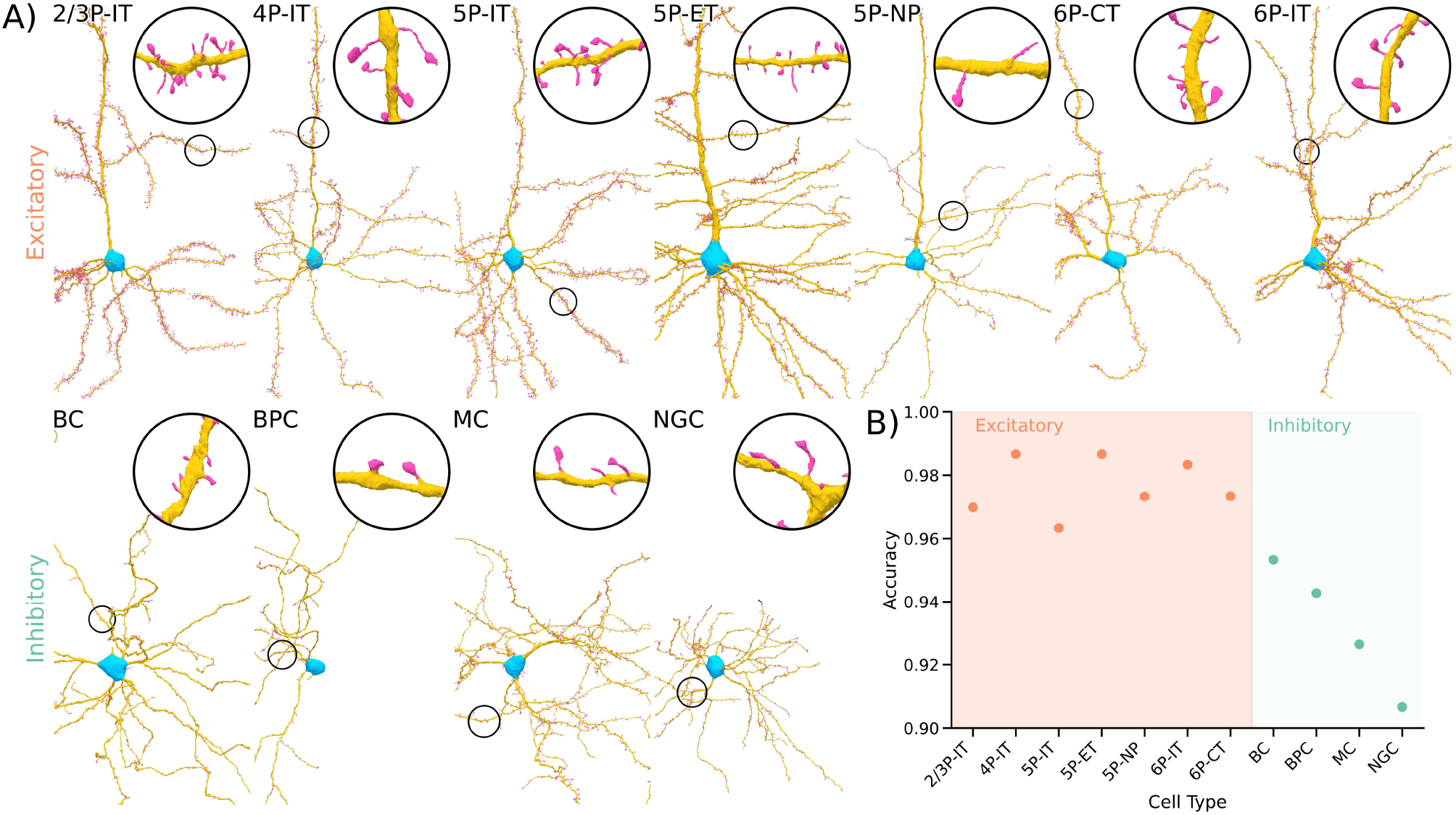
Depiction of model results on MICrONS and validation data. A) Example neurons from each of the main manually labeled cell types from Elabbady *et al*. [12]. Meshes are colored according to the predictions from the postsynaptic structure model, which was trained on point annotations only. Insets show zoom-ins on regions with putative spines. B) Validation accuracy on a set of synapses sampled from each cell type, for neurons in MICrONS not seen during training.

Because our previously described training data was collected in a manner that was not a uniform sampling of synapses and cell types, we wanted to ensure that our model was trustworthy when deployed “in the wild” on the MICrONS dataset. We independently collected a held-out validation set (all from previously unseen neurons at train time) by sampling 12 cells from each of the main cell types described in Schneider-Mizell *et al*. [29] and labeling 25 synapses at random on each of these cells. Classification accuracy on this sample of cell types and synapses remained high—in particular, synapses on excitatory neurons were classified with extremely high accuracy (weighted F1 score 0.977), while those on inhibitory neurons were more difficult but still fairly accurate (weighted F1 score 0.935) (Figure 3). We note that inhibitory dendrites appeared more variable in their fine morphology, with ambiguous spines or “thorns” that did not resemble processes on excitatory neurons, as well as larger variations in dendritic shaft thickness. These factors likely contributed to the lower accuracy on inhibitory neurons, and suggest future improvements to the model that are more robust to these variations (see section 3).

Nevertheless, this work represented the most accurate and comprehensive mapping of postsynaptic structures in a large electron microscopy volume to date (see Figure S4 for comparison to a previous approach, NEURD). While predictions are available for virtually all neurons in the dataset, for many of the subsequent analyses we focused on cells with proofread axons and manually labeled cell types to minimize the effect of errors from other automated pipelines on our analysis. We refer to this region of curated cells as the “column” following Schneider-Mizell *et al*. [29], as it is a representative sample of the cells present in a depth-spanning region in the center of the dataset.

### 2.4 Variation in spine targeting

Given this comprehensive information on postsynaptic structures produced by our pipeline, we first sought to understand variation in how cells receive inputs within and across cell types in MICrONS. Consistent with the long literature on mammalian neuron morphology, we found that excitatory neurons received more inputs onto spines than inhibitory cells (2159 vs 1043 on average, respectively) (Figure 4A). This distinction became even clearer when viewed as a proportion: 70.3% of input synapses onto excitatory cells in the column targeted spines, compared with only 19.8% for inhibitory neurons. Within these broad groups, there was further variability across cell types (Figure 4A). Excitatory intratelencephalic (IT) cells showed the largest proportion of their input synapses onto spines, in particular in Layer 2/3 and 4 (75.9% and 75.0%, respectively). Layer 5 extratelencephalic (ET), near-projecting (NP) and Layer 6 corticothalamic (CT) cells all showed slightly lower ratios of spine input (56.8%, 42.1%, and 62.5%, respectively). Among all of these cell types, there was also substantial within-type variation in the proportion of spine input. For instance, the range for Layer 5 ET cells was 32.6% to 73.3%. This is particularly remarkable given that the cells presented here are all adjacent in physical space, since they come from the same curated “column.” Among inhibitory cells, Martinotti cells (MC) had the highest average proportion of input onto spines (28.8%), followed by neurogliaform cells (NGC, 23.9%), bipolar cells (BPC, 17.5%), and with basket cells having the lowest proportion of spine input (BC, 11.3%). We note also that the Martinotti cells with the most spine input had a higher proportion than the lowest excitatory cells, showing that while spine input proportion is a good predictor of excitatory/inhibitory type, there is overlap in these distributions.

**Figure 4.**
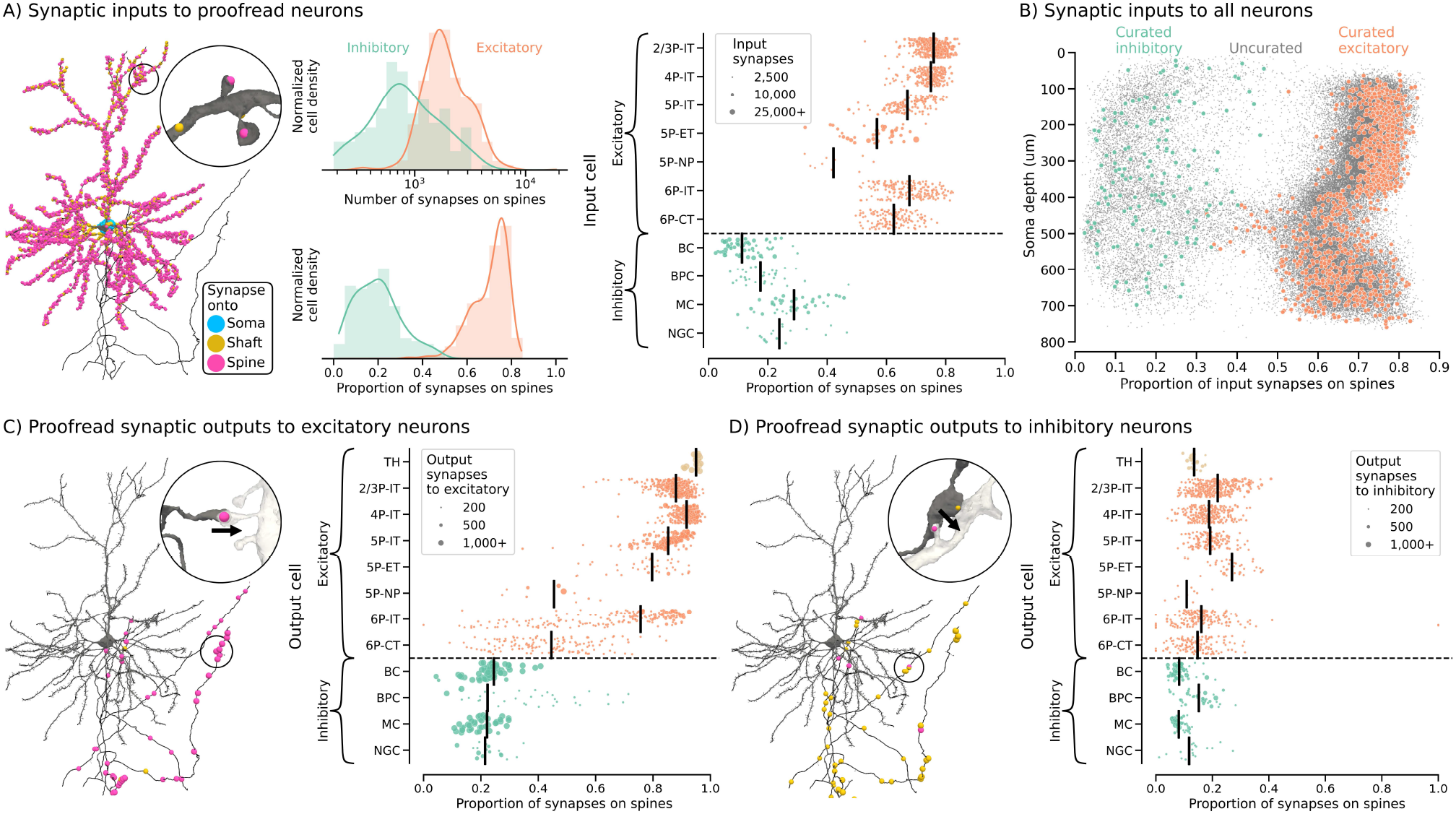
Description of target structures for inputs to and outputs from curated cells in MICrONS. A) Summary of all synaptic inputs onto curated cells. Left: diagram showing all input synapses onto an example cell as circles colored according to their predicted type. Middle top: histogram of the number of input synapses onto spines for excitatory and inhibitory neurons. Middle lower: histogram of the proportion of a neuron’s input synapses that are onto spines for excitatory and inhibitory neurons. Right: plot of each neuron’s proportion of input synapses onto spines. Neurons are sorted by cell type and within cell type by cortical depth (higher is closer to pia) on the y-axis. Black bars denote the weighted average proportion for each cell type, where weighting is by number of total input synapses for that neuron. Dots are also sized by the number of input synapses to that neuron. B) Comparison of proportion of input synapses onto spines between the curated neurons (large orange and green dots for excitatory and inhibitory neurons, respectively) and all other neurons in MICrONS. C) Synaptic outputs from cells with proofread axons onto putative excitatory neurons. Left: diagram showing all output synapses from an example cell onto excitatory neurons as circles colored according to predicted postsynaptic target type. Right: plot showing the proportion of output synapses onto putative excitatory cells which connect to a spine. Y-axis, black bars, and dots as in A) but with weighting by the total number of connections to excitatory neurons. D) Same as C), but for outputs onto putative inhibitory neurons.

The aforementioned analysis was focused on manually cell-typed neurons from the column, but we stress that this set of neurons was a dense, representative sample from all layers of cortex. Consequently, the patterns we observed mirror those in the rest of the dataset. We found that the distribution of proportion of spine input and the cortical depth of a cell’s soma was remarkably similar in our curated column to the entire dataset (Figure 4B). The slight shift towards lower spine input in excitatory cells can be explained by truncated dendrites, either due to being near the edge of the volume or the lack of proofreading for those cells.

We also aimed to study how cells’ outputs vary with regard to sub-cellular targeting patterns in this example of a local cortical circuit. In addition to the dendritic morphology information available from the EM reconstructions in the MICrONS data, we also had access to connectivity information. As expected, we observed that excitatory neurons predominantly targeted other excitatory neurons via spines (83.3%), while inhibitory neurons did so at a much lower rate (23.4%) (Figure 4C). In addition to the set of cells described previously, we also included the connectivity from a set of proofread thalamic axons entering the MICrONS visual cortex volume. These neurons had the highest preference for spine targets of any excitatory type (95.0%). More surprising was that when connecting to other excitatory cells, Layer 5 NP and Layer 6 CT cells targeted spines with much less preference than other excitatory cell groups (45.5% and 44.6%, respectively). Layer 5 NP and Layer 6 CT cells arrive late in development and from a shared lineage [30], raising the possibility that these cells share a developmental program which gives rise to this unusual feature of connectivity with other nearby excitatory cells.

Connection rates to inhibitory spines were less variable, reflecting the smaller proportion of spines on the dendrites of inhibitory cells overall. Excitatory cells were slightly more likely (19.1%) to target these inhibitory spines than other inhibitory cells were (8.2%). Among excitatory cells, Layer 5 ET cells were the most likely to contact inhibitory spines (26.9% of synapses from Layer 5 ET to inhibitory cells). Previous work in MICrONS observed that Layer 5 ET cells often connect to Martinotti cell spines [31]. We observed that this increased rate of spine targeting from Layer 5 ET cells was primarily driven by this effect (Figure S5). Further, no other excitatory cell type showed the same strong preference for Martinotti cell spines (Figure S5). Thus, our comprehensive map of postsynaptic targeting complements Bodor *et al*. [31] by showing the specificity of this connection mode (both for Martinotti and Layer 5 ET cells).

### 2.5 Multi-input spines

While spines on excitatory cells predominantly receive one excitatory input, multi-input spines are a well-documented phenomenon [32–36], and since our map included millions of spines, we sought to characterize their occurrence at scale. Such multiply-innervated spines most often have one excitatory and one inhibitory input [32–36], and have been documented as larger [33–35] and more stable [33] than singly-innervated spines, although it remains unclear whether this stability is the result of their larger size.

We wanted to examine how cells and cell types varied in the extent to which their spines displayed this pattern of multi-input innervation. To do so, we developed a scheme for finding putative multi-input spines from our mesh segmentation by finding connected components of the mesh which shared the “spine” label, and then found any groups of synapses which connected onto the same connected component (see subsection 4.7). Our manual review of 1,724 putative multi-input spines detected using this simple method found that 76% (95% CI: 74-78%) were true single spines with multiple synaptic inputs. This approach also detected other categories of postsynaptic structures at lower rates, namely 13% (95% CI: 11-14%) were “Y”-shaped spines which have two separate heads that each receive an input, and 6% (95% CI: 5-7%) were two separate spines which were fused in the automated segmentation or meshing. We developed a second classification strategy to separate these categories based on morphometry features for each putative spine connected component (see subsection 4.8). These features included the mean HKS features for each putative spine, as well as geometric features such as estimated volume, surface area, sphericity, and PCA-based descriptions of the principal extents of a spine component. A random forest classifier trained on these features was able to separate true unitary spines with multiple inputs from these other categories with 0.951 cross-validation accuracy. We note that our approach may miss some multi-input spines if the original synaptic detection classifier failed to detect one of the synapses on a multi-input spine or if our classifier failed to detect a spine, but we expect both of these failure modes to be rare based on the high accuracy of these models (precision of 96% and recall of 89% for the synapse detection model [9]).

We deployed this multi-input spine detection strategy across neurons in MICrONS. Of the 106.7 million putative spines detected by our model, 6.5 million were predicted to be unitary spines with multiple inputs. Multiple innervation of spines was rare but widespread across excitatory types (Figure 5A). Averaged across all excitatory cells, the rate of multi-spine innervation was 6.1%. This rate is comparable to quantifications in previous studies [8, 15, 32, 34, 37, 38], though we note this literature (drawn across multiple species, brain regions, and experimental modalities) cites rates that can vary from 3 to 26%. Spines on inhibitory cells more commonly received multiple inputs (Figure 5A), consistent with a previous study of spines near the somas of PV+ and SST+ interneurons in mouse primary visual cortex [39].

**Figure 5.**
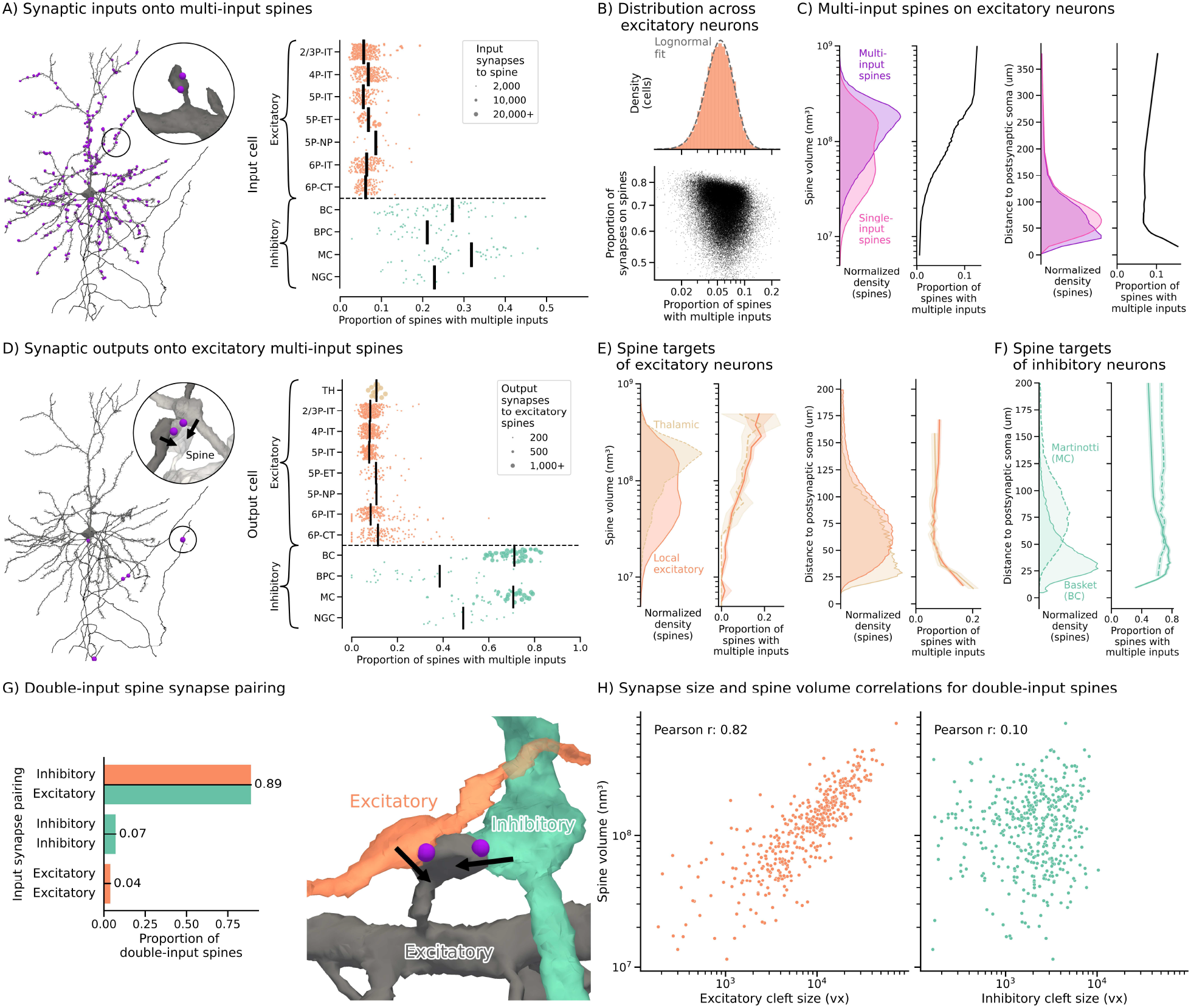
Multi-input spine targeting for proofread cells. A) Left: diagram of location of multi-input spines on an example neuron. Right: plot of neurons’s proportion of input spines with multiple inputs. Neurons sorted by cell type and within cell type by cortical depth (higher is closer to pia) on the y-axis. Bars show weighted average proportion by cell type; weighting by a neuron’s number of total input synapses onto spines. Dots sized by the same feature. B) Upper: distribution of excitatory neuron’s proportion of input spines with multiple inputs, shown with best lognormal fit. Lower: relationship between excitatory neurons’s proportion of input spines with multiple inputs and total number of input synapses onto spines. C) Left: histograms of spine volume for single- vs. multi-input spines, with line plot showing relative abundance of multi-input spines. Right: histograms of Euclidean distance from postsynaptic soma for single-vs. multi-input spines, line plot shows relative abundance of multi-input spines. D) Left: diagram of locations of outputs from an example neuron onto excitatory multi-input spines. Right: plot of each proofread neuron’s proportion of outputs onto multi-input spines, from its connections onto excitatory neuron spines. Y-axis, black bars, and dots as in A) but with weighting by total number of connections to excitatory neuron spines. E) Summary of sizes and locations of single- and multi-input spines as a function of presynaptic cell type. Left: Same as C) but as a function of whether the presynaptic cell is a thalamic or local excitatory cell. Right: Same as C) (right) but as a function of whether the presynaptic cell is a thalamic or local excitatory cell. F) Spatial distribution and multi-spine targeting rate as C) (right) for outputs from basket cells (BC) compared to Martinotti cells (MC). G) Left: proportion of curated double-input spines for which the presynaptic cells are inhibitory and excitatory, both inhibitory, or both excitatory. Right: example multi-input spine receiving one excitatory and one inhibitory input, which is by far the most common case. H) Examination of the correlation between spine size and cleft size for the excitatory (left) and inhibitory (right) connection onto manually verified double-input spines.

These previous works estimated spine multi-innervation rates from partial reconstructions of axons and/or dendrites and identification of their targets in EM, or sparse labeling of dendrites in light microscopy—to our knowledge no work has yet been able to examine variability in rates of multi-spine innervation across entire dendritic arbors and local circuits. We found that the mean rate of multi-spine innervation did not differ dramatically between excitatory types; however, there was substantial within-cell type variation. For instance, rates of multi-spine innervation on the 349 Layer 2/3 pyramidal cells in our curated column ranged from 2.2% to 19.3% (10th to 90th percentile was 3.5%-8.2%). This pattern of within-type heterogeneity also held across the rest of the dataset. Taking the population of excitatory neurons as a whole across all of MICrONS, the proportion of inputs onto spines was well described by a lognormal distribution (Figure 5B), although for individual excitatory cell types, some mixtures of lognormal distributions were observed (Figure S6).

We wondered what other factors of a cell’s morphology and collection of synaptic inputs could explain this variability. One hypothesis was that cells which received more excitation overall would have fewer multi-input spines, because inhibitory connections are known to co-innervate spines. As a proxy for the amount of excitatory input to a cell, we considered the proportion of a cell’s total input to spines—however, we found only a modest correlation between the total proportion of input to spines and multi-input spine proportions (Pearson’s *r* = −0.20) (Figure 5B). Similarly, within Layer 2/3 pyramidal neurons, we saw only modest correlations between a cell’s multi-input spine rate and cortical depth (*r* = 0.00), number of input synapses to shaft (*r* = 0.30), proportion of input synapses to shaft (*r* = 0.40), and mean input size (*r* = 0.46) (Figure S7).

Multi-input spines on excitatory cells were on average larger than single-input spines (Figure 5C), as previously described [32–36]. Further, we found a monotonically increasing relationship between the size of a spine and the probability that it was a multi-input spine (Figure 5C). Multi-input spines were also more prevalent both very close to the soma and in distal dendrites (Figure 5C). However, we found that a cell’s multi-input spine innervation rate was not strongly correlated with the average size of synaptic inputs onto its spines (Figure S7). The variance in multi-input spine rates for Layer 2/3 pyramidal cells is much larger than can be explained by a model in which the number and size of synapses on spines are kept constant, but multi-input spines are shuffled across neurons in the same cell type (Figure S8). Taken together, our census of tens of thousands of neurons in the MICrONS dataset revealed a remarkable within-cell-type variation in spine multi-innervation rates which is not fully explained by other simple morphological features measured here. This raises the question of what factors contribute to this diversity, and whether they can be explained by multiplicative processes which give rise to lognormal distributions, as described previously for spine sizes [40] and degree distributions and other network properties [41].

Since we also had access to proofread axons in the MICrONS volume, we were able to examine how cells targeted multi-input spines (Figure 5D). A previous immunohistochemistry study in the rat focused on the identity of the excitatory presynaptic connection to dually-innervated spines and found that almost all were innervated by a thalamocortical (VGLUT2+) axon [34], while almost all spines receiving input from a local excitatory (VGLUT1+) axon had only one input. Conversely, a more recent study in mouse visual cortex [37] using an expansion microscopy technique found less of a difference between thalamic and local excitatory rates of innervation to multi-input spines: across three labeled neurons, the authors found that 5/56 (8.2%) of thalamic inputs were onto multi-input spines, while 23/480 (4.6%) of local excitatory inputs were onto multi-input spines. Our results were more consistent with Balcioglu *et al*. [37] in that we observed only slight differences in rates of multi-input spine targeting: thalamic axons had a slightly higher rate (10.8%) of multi-input spine targeting compared to local excitatory axons (8.2%). Further, we saw that much of this difference could be explained by a shift in the size of the spines targeted by thalamic vs. local excitatory axons (Figure 5E). Thalamic axons tended to target larger spines than local excitatory axons, but the profile of multi-innervation rate as a function of size was similar between the two populations. Similarly, thalamic axons tended to target spines that were closer to the soma of the postsynaptic excitatory cell (Figure 5E), but the multi-innervation rate as a function of this distance was again similar. These observations suggest that the slight difference we and previous works observed may not require a distinct biological mechanism for multi-innervation by these two populations, but rather may be explained by differing statistics of spine size and target location by thalamic versus local excitatory axons.

In contrast to connections from excitatory axons, the majority of excitatory spines which were contacted by inhibitory neurons had at least one other input (70.2%). This phenomenon was most dramatic for basket cell (BC) and Martinotti cell (MC) types (71.2% and 70.7%, respectively), which together make up the majority of inhibitory synapses and align well with previous electron microscopy measurements of sparsely labeled cells in the rat [34]. We wondered whether there was any patterning to the synapses made by inhibitory neurons onto singly-vs. multiply-innervated spines. We compared connections from basket cells, which are known for their targeting of the perisomatic region of other neurons, to Martinotti cells, which are known to target excitatory neurons more distally. We observed this expected difference in the distribution of distances to the postsynaptic soma when comparing Martinotti and basket cell outputs (Figure 5F). However, we also found that basket cells often targeted single-input spines on or very close to the soma (Figure 5F). Multi-input spine targeting from basket cells quickly increased to a peak of around 80% at approximately 40 microns from the soma, before steadily decreasing again. In contrast, Martinotti cells consistently targeted multi-input spines at a similar rate across all distances from the soma. This suggests that the spines targeted by basket cells close to the soma may be a distinct population. This observation complements previous studies in other brain regions which have shown that the distribution of organelles and receptors within these somatic spines is different [42, 43].

Finally, we also examined cases where not just one, but both inputs to a multi-input spine were known. We selected spines where both input axons had been proofread and connected back to a soma with a known cell type. Further, we manually reviewed these synapses to select only cases where both axons were synapsing onto the same single-headed spine, as opposed to being in a Y-configuration or other failure mode for the classifier (see section 3). In the vast majority of cases, excitatory (including thalamic) synapses were paired with an inhibitory synapse (Figure 5G). We further examined the size of the presynaptic clefts (determined by an automated machine learning classifier as described previously [9]) in relation to the size of the spine (Figure 5H). We observed that, consistent with single-input spines, there was a strong relationship between the size of the spine and the size of the excitatory cleft (Pearson’s *r* = 0.82). Further, this relationship closely matched that of other putative single input spines (Figure S9). In contrast, there was a very weak relationship between the size of the spine and that of the inhibitory cleft (Pearson’s *r* = 0.10). This lack of a strong relationship between the inhibitory cleft size and spine volume could be explained by the dynamic nature of these connections [33], as well as the smaller relative size of the inhibitory connections overall.

### 2.6 Transfer to H01 connectome

We wanted to test the extent to which the methods described here could be applied out of the box to other connectome datasets to enable similar analyses. In several other datasets, we observed HKS features along neuronal arbors that qualitatively mirrored what we observed in MICrONS (Figure S1). To quantify the generalizability of our approach, we applied the HKS pipeline and postsynaptic structure classifier to pyramidal neurons in the H01 connectome dataset [11], a reconstruction of a cubic millimeter of human temporal cortex. We randomly selected 80 pyramidal neurons, and manually labeled 25 synapses on each using the same criteria as described in subsection 4.2. We then computed the same HKS features without changing any parameters, and applied the postsynaptic compartment classifier fit on the MICrONS dataset “zero-shot” without any retraining of the model.

This model maintained high accuracy on this dataset, with a weighted F1 score of 0.949 (compared to on pyramidal neurons in MICrONS, Figure 6A). In contrast with MICrONS, predicted somatic regions were enlarged relative to the manually labeled sites (Figure 6B), likely due to differences in the absolute biological scale of neurons in two datasets. Similarly, in the H01 dataset we observed more true shaft synapses labeled as spines than the converse (Figure 6B), a trend we observed only slightly in MICrONS (Figure S3). This difference is likely caused by a difference in scale, as misclassifications were more often observed on larger spines (Figure 6D, inset 2). Though spines on human pyramidal neurons do largely overlap in size distribution with those of mice, they are larger on average, and the largest human spine heads have more than twice the volume of the largest mouse spine heads [2]. We also observed some misclassifications at artificial borders in the mesh caused by edge effects from gaps across missing slices which had subsequently been corrected in the segmentation, though these issues were localized (Figure 6D, inset 3). Despite these issues, these results demonstrate strong transfer performance for a model which only had access to labels from one dataset. Further, the model was sufficiently calibrated (Figure 6C), meaning that the confidence of the model tracked with the empirical probability that a given prediction was correct. This suggests that future labeling for fine-tuning a model to a new dataset could likely be achieved by selecting uncertain regions for labeling.

**Figure 6.**
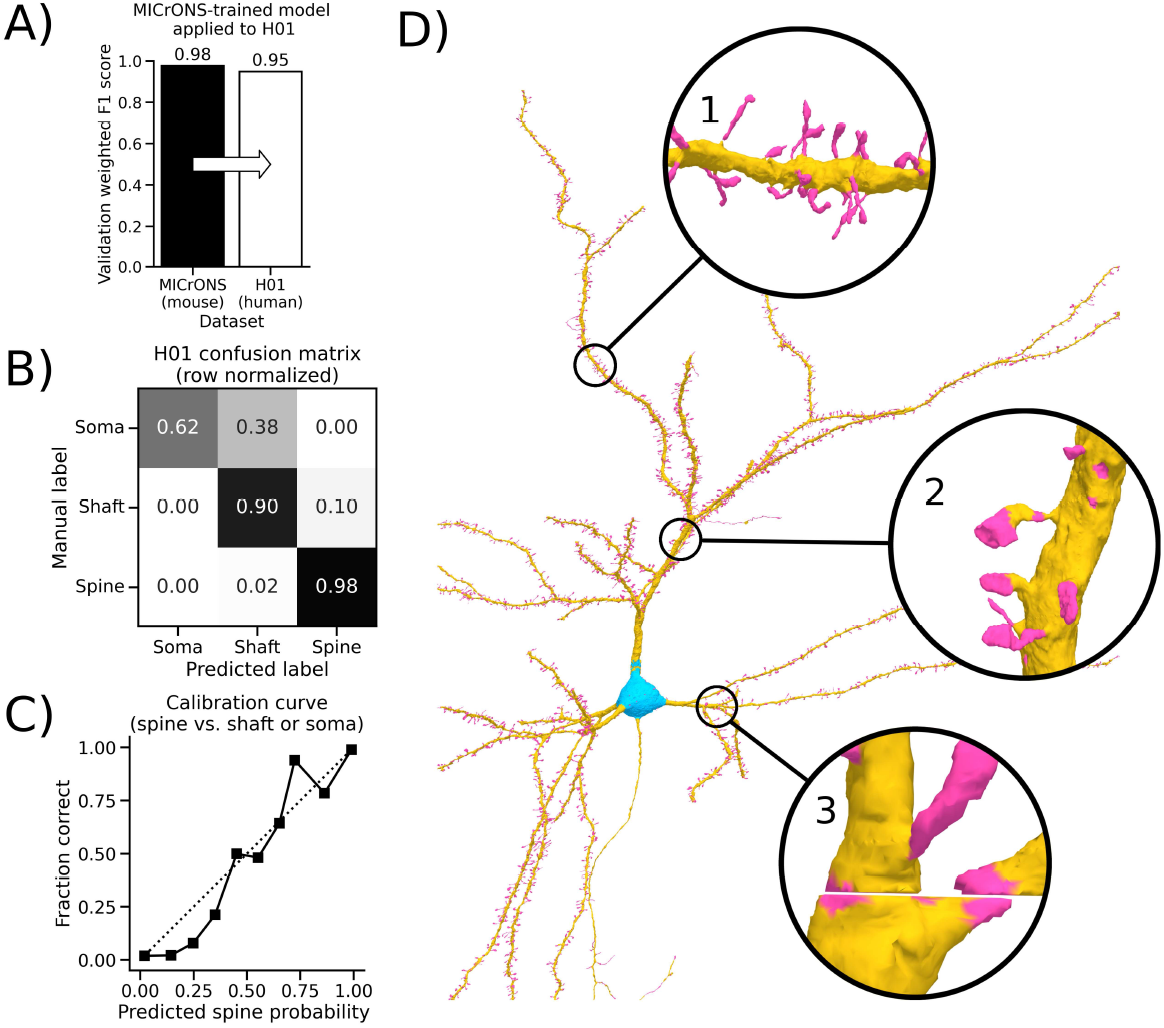
Application of the HKS method to the H01 connectome. A) We performed zero-shot transfer of our HKS-based postsynaptic structure classifier to the H01 connectome, a reconstruction of a piece of human temporal cortex. On independently collected validation labels, the model still performed well (0.949 weighted F1 score). B) Confusion matrix for the same experiment described in A). The model performed worst on somatic regions which were more often misclassified as shafts, likely due to differences in absolute scale between the two datasets. C) Binary calibration curve showing relationship between the model posterior (probability of a sample being predicted spine) and the empirical proportion of samples which were spines in the validation set. D) Model predictions shown on the mesh for an example neuron from the H01 dataset. Inset 1 - example of correctly segmented spine and shaft regions; 2 - example of very large spines which were partially misclassified as shafts; 3 - example of a missing slice artifact which caused an edge effect where some parts of the shaft at the border were classified as spines.

## 3 Discussion

### 3.1 Contribution

We present a method for postsynaptic structure prediction based on the mesh of a neuron, a ubiquitous structure in modern EM datasets that is often used to represent neurons for visualization. We showed that this classifier worked well across many cells from different cell types. Further, we developed a computational pipeline that could construct these mesh-based features robustly in the commercial cloud at a moderate computational cost.

To demonstrate the utility of our approach, we applied it to the MICrONS dataset, and described some of the initial results from this map of over 208.6 million postsynaptic sites. We characterized the variability in spine input and output targeting rates across and within cell types. As expected, excitatory neurons received more of their inputs onto spines. Similarly, most excitatory types predominantly targeted spines when connecting to other excitatory cells, with the notable exceptions of Layer 5 near-projecting (NP) and Layer 6 corticothalamic (CT) cells, which often targeted shafts of other excitatory cells. All inhibitory cell types targeted spines at lower, though still substantial rates when connecting to excitatory cells. Within cell types, we generally observed substantial variability in spine targeting rates (both for input and output synapses).

Additionally, we developed a method for detecting multi-input spines, and again used this method to describe variability of multi-input spine innervation across MICrONS. We found that rates of multi-input spine innervation varied widely within excitatory neurons (even of the same type) and that this variability was well characterized by a lognormal distribution. Given the large number of spines in MICrONS, we were also able to examine how a cell’s rate of spine multi-innervation varied with factors such as spine size, distance from the soma of the postsynaptic cell, and the identity of the presynaptic cell type. We observed little correlation with these factors, which may indicate that dynamic processes not observed in this static connectome contribute to this variability.

While we presented some initial analysis of this census of subcellular targeting, we expect that the greatest use for these annotations will be for future researchers interested in querying synaptic connectivity and morphology in MICrONS. The synaptic annotations are available for the MICrONS dataset for anyone to access via CAVE [44] (table name `synapse_target_predictions_ssa_v2`).

Beyond MICrONS, we also applied the feature generation pipeline to example structures from a variety of recent connectomics reconstructions, including from a mouse hippocampus [45], a songbird basal ganglia [46], and a fly brain [47] (Figure S1). These proofs of concept demonstrate that the pipeline is easy to run on any reconstruction with meshes, and that the features can likely be used to quantify a variety of neuroanatomical structures. For instance, the fly neuron shows a clear feature separation (even for the one HKS timescale shown) at “twigs,” small input structures on dendrites. In the mammalian datasets, axonal boutons also differ from the non-bouton portions of the axon across a variety of cell types and datasets. We also demonstrated that our postsynaptic structure classifier trained on the MICrONS dataset could be applied out of the box to the H01 connectome [11] with good performance, further indicating that the methods developed here can be used on other datasets with minimal adaptation. To facilitate the future reuse of these tools, we have created a website which includes links to the code for HKS feature generation and cloud deployment, the postsynaptic structure model, as well as a web-based demo of these tools to enable researchers to quickly try these techniques on their own data ^5^.

### 3.2 Limitations and future work

Meshes are convenient representations of morphology because they are compact relative to the size of the segmentation while preserving a cell’s exterior geometry. They are also ubiquitous in modern connectome datasets, making it easy to apply mesh-based models to new data. However, the reliance on surface geometry and connectivity also means that the model is susceptible to errors in the mesh, which are common in connectome datasets as mesh generation pipelines have mainly been optimized for speed, storage, and visual fidelity. We observed subtle issues with the model where the mesh is completely discontinuous across gaps (Figure 6). While the robust Laplacian approach we leverage in HKS computation can help alleviate issues from some mesh degeneracies [26], complete discontinuity causes long-timescale HKS features to be distorted by an artificial boundary. Future work should explore whether augmentation (i.e., artificially injecting these gaps into training data) or recently described frameworks for dealing with topologically broken objects [48] could help mitigate this issue.

The application of our classifier to the H01 dataset suggested that the model has some sensitivity to scale (e.g., dendritic shaft diameter or spine size), which is natural given the nature of HKS features which do depend on the absolute size of objects [49]. Similarly, we observed that some misclassifications in MICrONS also appeared related to absolute scale; i.e., some narrowings of dendritic shafts could be misclassified as spines (though, note that these often are devoid of synapses and thus not as common in the synapse-based training data and subsequent analysis). While this scale dependence can be a helpful prior (e.g., spines are usually smaller than shafts), it can also lead to generalization issues between datasets and organisms. Future work should investigate this tradeoff by training models with augmented data where this scale is varied over some range during training, or explore mathematical approaches to reducing the scale dependence of the HKS features [49]. Training models with data from multiple connectome samples could also be a sufficient and natural way to create a more generalizable model.

We also note that while the HKS features clearly contain much of the information required to classify postsynaptic structures, future models may benefit from higher-order geometric context to further improve performance or to be applicable to other tasks. Many works have built upon the HKS and related spectral descriptors, including by building secondary features from collections of HKS features on a shape [50] or by learning task-specific spectral filters [51]. These approaches were in many ways precursors to modern geometric deep learning approaches [52] such as graph neural networks. However, even with these approaches, state-of-the-art performance is often still achieved by leveraging the HKS or related spectral descriptors as base features [53, 54]. This suggests that our proposed pipeline for computing and storing HKS representations could be augmented by these higher-order learning approaches to further improve performance or to enable other morphological learning tasks, such as segmenting axonal boutons or classifying cell types.

## 4 Methods

### 4.1 MICrONS dataset

The MICrONS structural dataset is an electron microscopy volume of visual cortex from an adult mouse (P87). We refer to The MICrONS Consortium *et al*. [9] for extensive details about how this volume was collected, imaged, and reconstructed. The dataset is continuously undergoing proofreading and annotation; for this work, we used version 1412 from CAVE [44].

### 4.2 Manual labeling of postsynaptic targets

Initial manual labeling of synaptic target sites was performed as part of the Virtual Observatory of the Cortex (VORTEX) program, and the labels are available from CAVE [44] table `vortex_compartment_targets`. Synapses were selected under one of two regimes: 1) all automatically detected synapses that were the outputs of inhibitory cells (25 cells, ~90,000 synapses), 2) all automatically detected synapses that were the inputs onto excitatory neurons (7 cells, ~30,000 synapses). A further ~4000 synapses were selected from all automatically detected synaptic outputs of layer 5 extratelencephalic (L5-ET) pyramidal cells, and labeled as part of analysis for [31].

Independent of how the synapses were selected, every synapse was manually inspected in Neuroglancer [55] by expert annotators. The presynaptic and postsynaptic meshes on either side of the synapse were rendered, and the annotator tagged the postsynaptic compartment as one of “spine”, “shaft”, “soma”, “soma spine”, “orphan”, or “other”. The annotator also reviewed the imagery of the synaptic cleft to decide ambiguous cases, for example where an axonal bouton contacted both spine head and dendritic shaft. To be labeled a spine, the postsynaptic compartment beneath the synapse needed to show distinct narrowing and physical separation from the dendritic shaft. “Stubby” spines, where the spine neck was indistinguishable from spine head, were labeled spine. Synapses onto the neck of a spine, rather than the spine head, were labeled spine. Synapses onto the dendritic shaft at the base of a spine neck were labeled “shaft”. The “soma” label was reserved for any synapses onto the soma of the cell, except for those on spines directly protruding from the soma. These had their own label of “soma spine.” The distinction between “soma” and “shaft” was the pronounced tapering of the dendritic branches from the soma. Some broad apical trunks that showed no tapering were treated as continuous with soma, until the first branching or narrowing was observed. If the postsynaptic compartment was a small, disconnected object, the synapse was labeled “orphan”. These were often, but not always, spine heads whose segmentation was disconnected from the dendrite. The label “other” was a catch-all for synapses that do not match other labels, for example synapses onto astrocytes or onto neuronal axons. Synapses that were found to be errors of automatic synapse detection were removed from further analysis.

### 4.3 Pipeline for heat kernel signatures

Below, we detail the steps of our proposed pipeline for computing heat kernel signatures.

#### Mesh download

Meshes were originally created from the underlying segmentation using the fast marching cubes algorithm [56] in zMesh [57], followed by quadric mesh simplification [25] as described in Macrina *et al*. [58]. We accessed these meshes using CloudVolume [59].

#### Mesh simplification

Meshes were simplified further again using the quadric mesh simplification algorithm [25], as implemented in fast-simplification [60]. We simplified the meshes provided by the above pipeline by 70% target decimation, and with the “aggression” threshold set to 7. We chose these parameters based on the performance of the resulting HKS algorithm run on the simplified mesh, choosing the highest degree of simplification which did not lower the performance of the resulting classifier on the postsynaptic structure labeling task. We note that the parameter choices for this part of the pipeline should depend on the length scales and resolution of the meshes that one is starting from. For Figure S1 only, we set the simplification parameters by setting the target vertex density (mesh vertices per unit surface area) to match that of the simplified meshes in MICrONS, which was approximately 5 × 10^−5^ vertices per nm^2^.

#### Overlapping mesh splitting

To make the eigendecomposition of the Laplacian scalable, we first subdivide the mesh into smaller pieces, which can optionally be processed in parallel. We note here that much like when estimating a spectrogram, there is a tradeoff between the domain (here, the size of the mesh) and the frequencies which can be accurately estimated (here, the timescales of the heat kernel signature features). Subdividing the mesh reduces our ability to accurately represent heat diffusion over very long timescales; however, we empirically observed that this did not affect our ability to perform spine detection, which requires only local information on the order of 5-10 microns.

This mesh subdivision proceeded in two stages: the first to determine the mesh chunks, and the second to add overlap so that the re-stitched computation on the mesh was smooth without boundary artifacts. To determine the mesh splits, we constructed a graph Laplacian based on the connectivity of the mesh, and performed an approximate graph bisection by splitting the mesh based on whether a vertex’s value of the second smallest eigenvector was positive or negative [61]. This process recursed on the resulting submeshes until all child submeshes were smaller than 20,000 vertices; this submesh was defined as the “core”.

During the second phase, these core submeshes were grown by some amount so that there was overlap between grown submeshes. This overlap minimized edge effects, which would otherwise create sharp transitions at the boundaries between submeshes. We grew each core submesh by 20 microns or the 60,000-nearest vertices, whichever was smaller. This avoided automatically including the large meshes with many vertices (usually falsely merged astrocytes) into calculations, as the cost of the decompositions scales with the size of the submesh. During this stage we kept track of which vertices in the grown submesh were part of the original core; the subsequent computed HKS values for these core vertices would be kept while those for the grown region of overlap would be discarded, as those vertices would be in the core set for another submesh.

#### Construction of the robust Laplacian

Numerical schemes for computing the HKS and related spectral descriptions of mesh geometry often suffer from issues at non-manifold regions (e.g., holes) of the mesh. These imperfections are common in modern meshing pipelines used in connectomics, mainly because they were optimized for speed and hierarchical chunked parallelization such that they could scale to the size of modern datasets and facilitate real-time meshing for proofreading. One approach to alleviate this issue is to attempt to post-hoc “fix” the mesh–however, in practice we found that many of these tools would have issues at thin processes and often would further disconnect pieces of the mesh. Instead, we opted to use the robust mesh Laplacian of Sharp & Crane [26]. Rather than attempt to modify the mesh itself, this approach simply modifies the mesh’s corresponding discrete Laplacian matrix. This approach was extremely fast, led to stable heat kernel solutions, and was easy to implement using the existing code for this approach. We set the mollification factor as described in Sharp & Crane [26] to 1 × 10^−5^. We note that many alternative schemes for determining the initial chunking of the mesh could be used; for instance, spatial chunks aligned to an octree would have certain advantages for distributed processing. We leave these improvements to future work.

#### Eigendecomposition via the band-by-band algorithm

Given the mesh’s discrete Laplacian matrix, we next needed to compute the partial eigendecomposition of some number of smallest eigenvalues. The Laplacian matrix of a mesh is extremely sparse, making sparse iterative eigensolvers such as ARPACK an attractive solution. However, these approaches are best at solving for the largest or smallest eigenvalues of a matrix, and often slow down greatly if one needs to compute more than a small number of eigenpairs. We found that to get resolution with the heat kernel down to the scale of spines, eigendecompositions might need on the order of hundreds of eigenpairs. We leveraged the approach described by Vallet & Lévy [27], which was developed to solve exactly this problem. This so-called “band-by-band” approach computes eigenpairs in bands around some value using a well-known “shift-and-invert” trick to speed computation of eigenvalues around some set point. Following the original authors, we started with a band centered at 0, and computed eigenvalues in bands of 50 eigenpairs at a time. After each band, we shifted the band up by 40% of the most recent band’s width (difference between maximum and minimum eigenvalues), and repeated the process. We ensured that the resulting bands overlapped and recomputed for a smaller change in band if not. We terminated the eigendecomposition after reaching a maximum eigenvalue *λ*_*max*_ = 1 × 10^−5^. We note that this value should depend on the length scales in the Laplacian matrix, which in turn depend on the units of the mesh. Our meshes were in units of nanometers.

#### Heat kernel signature computation

Given the partial eigendecomposition of each submesh, we next computed the truncated approximation of the heat kernel signatures for each point in the submesh. Following Sun *et al*. [24], the heat kernel signature for a given timescale *t* could be approximated as

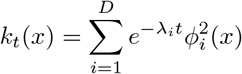

where *D* is the number of computed eigenpairs for that submesh, and *λ*_*i*_ and *ϕ*_*i*_ are the *i*th eigenvalues and eigenvectors of the robust mesh Laplacian, respectively. Given the eigendecomposition, this computation is extremely fast, and could be repeated for any specified timescale. Again following Sun *et al*. [24], we used logarithmically spaced timescales between *t*_*min*_ and *t*_*max*_. *t*_*min*_ and *t*_*max*_ were set to 5 × 10^4^ and 2 × 10^7^. We set the number of timesteps between these values to 32—above this number we did not notice an appreciable change in performance.

#### Agglomeration of mesh features to local supervoxels

The approach described thus far yields a HKS feature vector for each vertex in a given mesh or submesh—for a neuron, this can amount to several million vertices. We developed a simple lossy compression scheme for reducing the size of storing these mesh-based features for later reuse, which we found valuable for the ability to update a classifier while leaving the computed HKS features which required the majority of the computation for this pipeline. Our approach used a simple connectivity-constrained agglomerative clustering (Ward’s method, though any linkage criterion could be used) as implemented in scikit-learn [28] to find small clusters of mesh vertices which had relatively similar HKS feature vectors. The only hyperparameter for this method was the distance threshold for where to cut the agglomerative clustering. We set this distance threshold to 3. The connectivity-constrained agglomeration also ensured that these clusters consisted only of single connected components of vertices. To compress features using this scheme, we saved the mean HKS feature for each cluster and the mapping from mesh vertex to cluster identity. Both of these arrays were compressed into a single file using NumPy [62] and gzip compression.

#### Auxiliary features

In addition to the HKS features, we also found it helpful to include distance to the neuron’s nucleus center (computed as described previously [9]) as an additional feature for each mesh vertex (or, using the compressed scheme above, the distance to the nucleus center from the centroid of each compressed mesh region).

### 4.4 Pipeline deployment via commercial cloud

We used the commercial cloud (Google Kubernetes Engine) to deploy the computational scheme described above at scale. We stress that the core code described above still operates on a single mesh at a time, and is not in any way locked to a particular cloud provider or method of deployment. We used “c2d-highmem-32” machines on Google Kubernetes Engine, each of which provided 32 vCPUs and 256 GB of RAM per node. At peak operation, we used 32 such machines, and each machine had 32 independent processes all running in parallel to download meshes and run the aforementioned pipeline. Results were saved using the compressed agglomeration scheme (see subsection 4.3) to a cloud object store (Google Cloud Storage) for later retrieval. More details about our specific deployment are available at github.com/bdpedigo/cloud-mesh.

### 4.5 Classifier

We used a random forest classifier [63] as implemented in scikit-learn [28] to classify postsynaptic structures based on the labels and features described above (subsection 4.2 and subsection 4.3, respectively). The “soma” and “soma spine” labels were combined as “soma”. “Orphan” and “other” were dropped for the purposes of training and evaluation. The hyperparameters for the random forest used the default values in `scikit-learn` except for the number of trees (`n_estimators =` 500) and the maximum depth of each tree (`max_depth =` 15).

### 4.6 Validation set

To ensure our classifier performance on our training data would generalize to the rest of MICrONS, we collected validation labels sampled broadly across the dataset and various cell types. We randomly selected 12 neurons from each of the major cell types considered in this work (see subsection 4.1). Cell type classifications for this purpose came from Elabbady *et al*. [12]. Inhibitory cell classes were further stratified across the soma depth of the cell to ensure sampling across all cortical layers. We manually reviewed these cells and excluded any for which we disagreed with the automated cell type classification or which had very incomplete dendritic reconstructions, resampling a new neuron from that class if a cell was excluded. From each of the selected neurons, we randomly sampled 25 automatically detected synapses which were onto that neuron. A human annotator then reviewed each of these synaptic sites in Neuroglancer [55] and labeled the postsynaptic structure as spine, shaft, soma, or other (e.g., axon initial segment, etc.). “Other” labels were ignored for the purposes of evaluating classifier performance. In total, this validation dataset consisted of 300 synapses per cell type, or 3,300 synapses overall.

### 4.7 Spine connected components and morphometric features

After a given neuron’s mesh was run through the HKS pipeline and the subsequent classifier, we extracted submeshes that contained a connected component of “spine”-labeled vertices. This was accomplished by dropping all mesh edges which connected vertices with different predicted labels and then running the connected components algorithm in SciPy [64] on the resulting graph. We then extracted the induced submesh associated with the vertices of each of these connected components; this was considered a putative spine. We mapped synapses to the spine-labeled connected components which they connected to in order to find cases of putative multi-input spines (see subsection 2.5).

We also estimated the volume and other morphometric properties of each putative spine using the extracted submeshes. After extraction from the neuron mesh, each spine submesh was not watertight (e.g., at minimum there would be a hole where the spine connects to the dendritic shaft), making simple methods for volume estimation on meshes unstable. Rather than attempt to “fix” each of these meshes, we randomly sampled points in a tight bounding box around each submesh, and estimated which points were interior or exterior to the spine using the fast winding number approach of Barill *et al*. [65]. This method is a robust way of estimating whether a point lies inside a given mesh even in the presence of holes. The induced submesh and the corresponding interior point cloud were used to compute a variety of features:

- The surface area of the spine (*A*) was computed directly from the mesh faces.
- The approximate volume (*V*) of the spine was then computed using the ratio of interior to exterior points multiplied by the volume of the tight bounding box.
- The sphericity of the spine was computed as 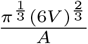
- PCA was computed on the interior point cloud, and the lengths of the three principal axes (*p*_1_, *p*_2_, *p*_3_) were estimated using the singular values of the mean-centered point cloud.
- The second smallest eigenvalue (or the so called “Fiedler value”) of the graph Laplacian of the submesh was computed as a measure of how well connected the mesh was.
- The mean and maximum distance to the boundary of the spine mesh were computed from the population of interior sampled points.
- The mean values of each HKS feature across all vertices in the spine mesh were also computed.

### 4.8 Unitary spine classifier

Using the features described in subsection 4.7, we additionally trained a random forest classifier [28, 63] to classify whether a given putative spine connected component with more than one synaptic input was indeed a true single spine (i.e., a single-headed spine connected to the dendritic shaft) or a false positive (e.g., a Y-shaped spine with multiple distinct heads or two or more spines merged together). Our training data for this task came from manually reviewing 1,724 putative multi-input spines sampled from MICrONS. We sampled these synapses using a similar strategy to subsection 4.6, selecting 10 neurons from each of the major excitatory cell types and randomly sampling a maximum of 25 putative multi-input spines from each neuron (fewer if there were not 25 putative multi-input spines on that cell). A human annotator labeled each of these sites as whether it was a distinct unitary spine with a single head as opposed to the other categories described above or another structure or misclassification. The random forest classifier used the default hyperparameters in `scikit-learn` except for the number of trees (`n_estimators =` 500) and the maximum depth of each tree (`max_depth =` 15).

### 4.9 Software

Software used in this work includes: NumPy [62], SciPy [64], scikit-learn [28], PyVista [66], CloudVolume [59], cmasher [67], point-cloud-utils [68], and gpytoolbox [69].

## 5 Supplemental Figures

**Figure S1:**
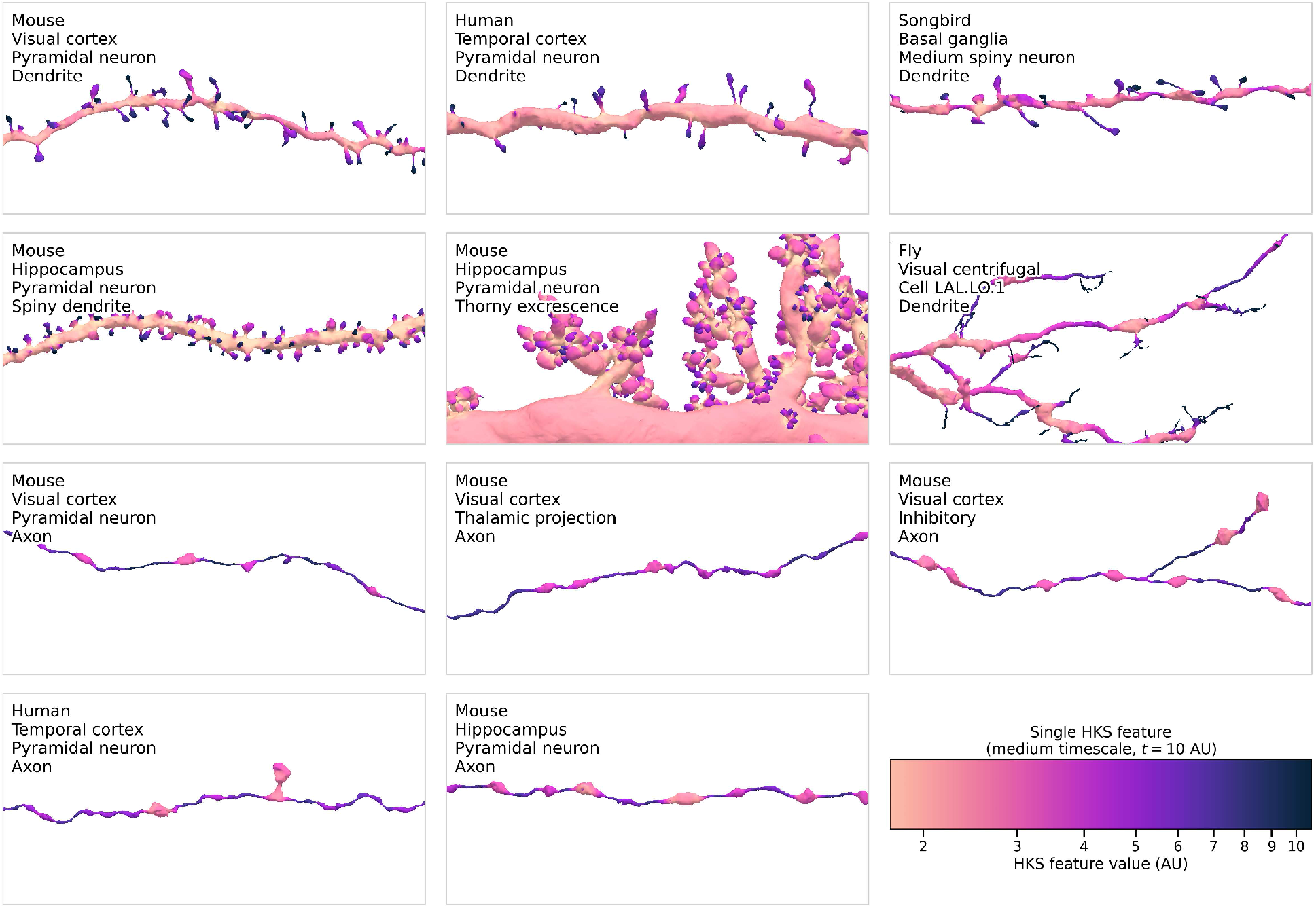
One HKS feature (i.e., a specific timescale for diffusion) shown across selected objects in MICrONS [9], H01[11], songbird basal ganglia[46], mouse hippocampus[45], and Flywire FAFB [47]. All meshes were simplified to have approximately the same vertex density (see subsection 4.3 for details), and the HKS features uses the same scale across figures. Note that variation in the HKS feature remains largely similar across homologous structures; spines have a higher HKS and boutons have a lower HKS than their surrounding.

**Figure S2:**
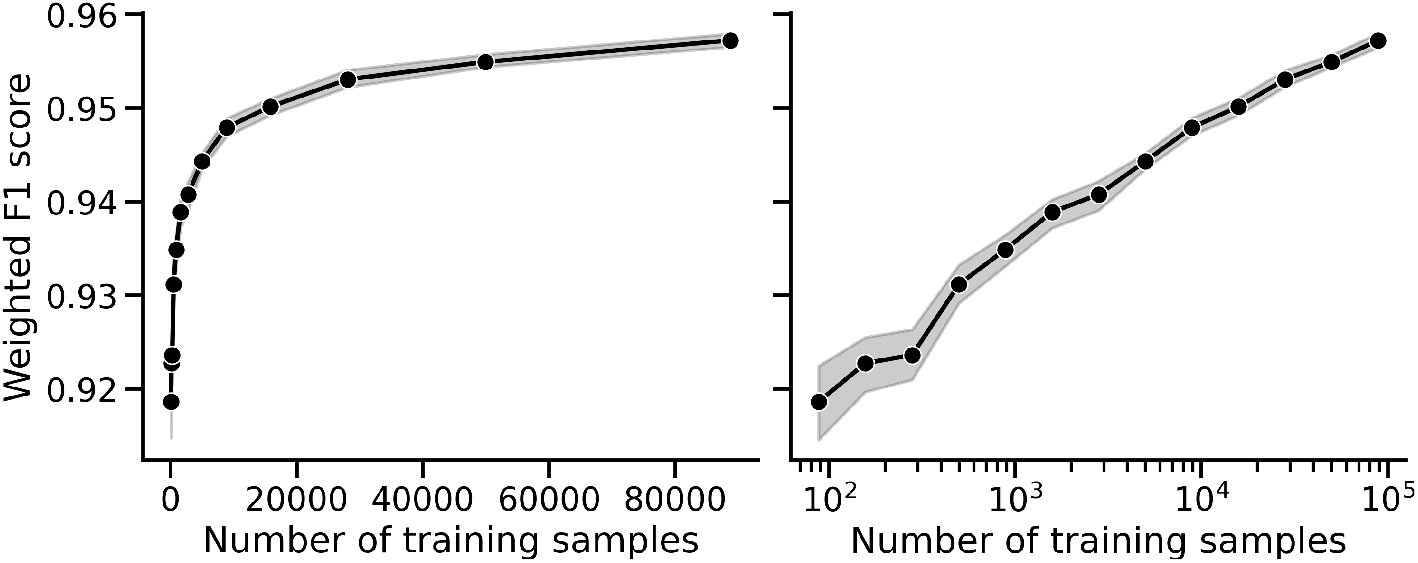
Classifier performance (weighted F1 score) as a function of the size of the training dataset. Same data is shown but on a linear (left) and logarithmic (right) x-axis. Each point shows mean and 95% confidence intervals across 10 random seeds; 20% of training data was held out for evaluation for each experiment.

**Figure S3:**
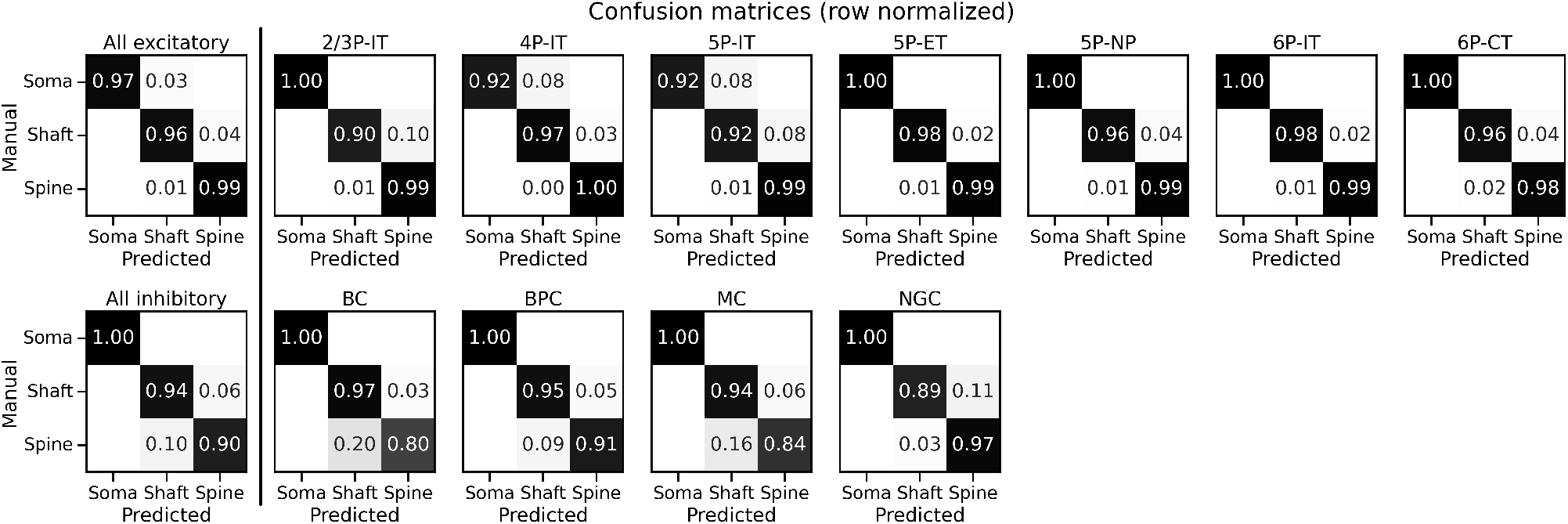
Confusion matrices within each cell type for the validation experiment described in Figure 3. Matrices are normalized by the count of the manual label in each row. Each cell type’s validation data consisted of 300 total randomly sampled synapses, evenly distributed across 12 randomly sampled neurons of that type.

**Figure S4:**
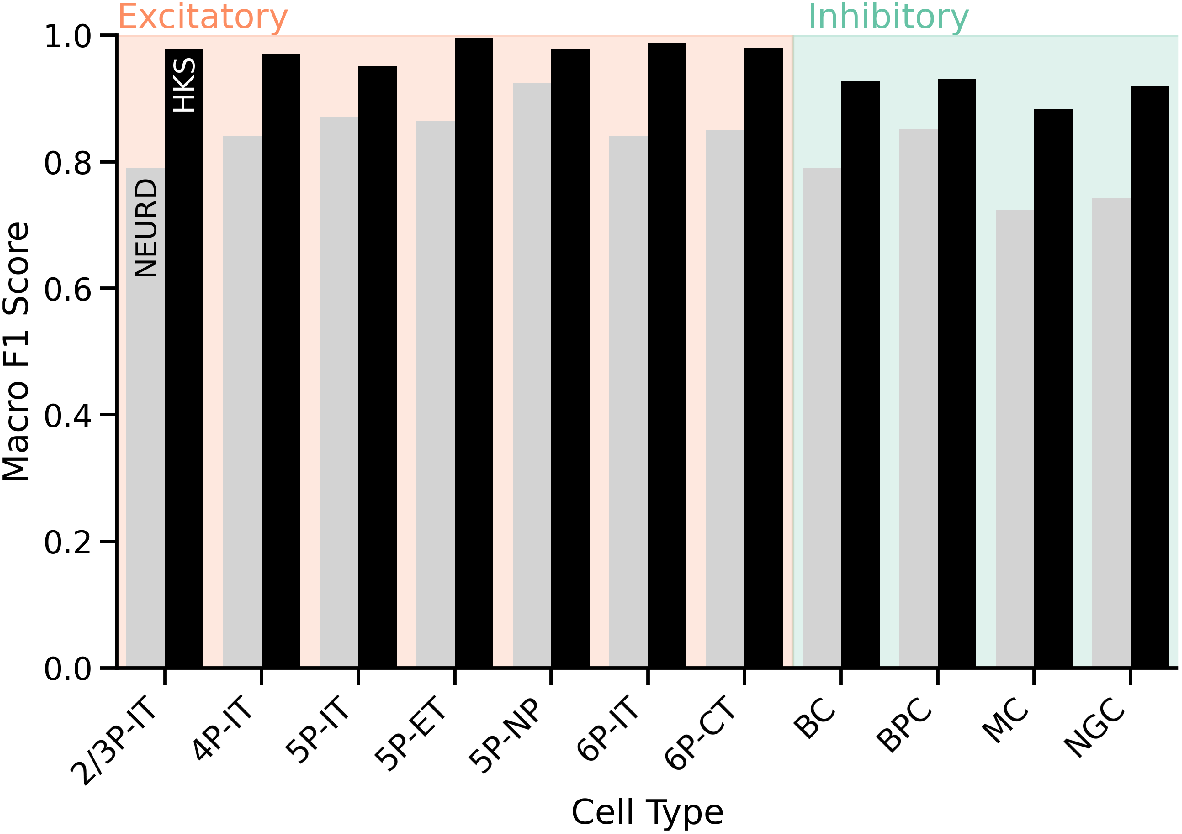
Comparison of classifier performance to NEURD [14], a previously described method for postsynaptic target classification which was applied broadly across MICrONS. Models were evaluated on the validation data collected for Figure 3. We selected the subset of samples for which model predictions from both methods were available. NEURD predictions were accessed from the CAVE table “synapse_target_structure” For this validation set (which neither model saw for training) our proposed HKS-based classifier was more accurate (weighted average F1 score 0.842 for NEURD, 0.963 for our proposed HKS scheme). This trend held across all cell types. We note that NEURD classifications were simplified to only the spine, shaft, and soma labels, as NEURD attempts to split spines into head and neck where possible.

**Figure S5:**
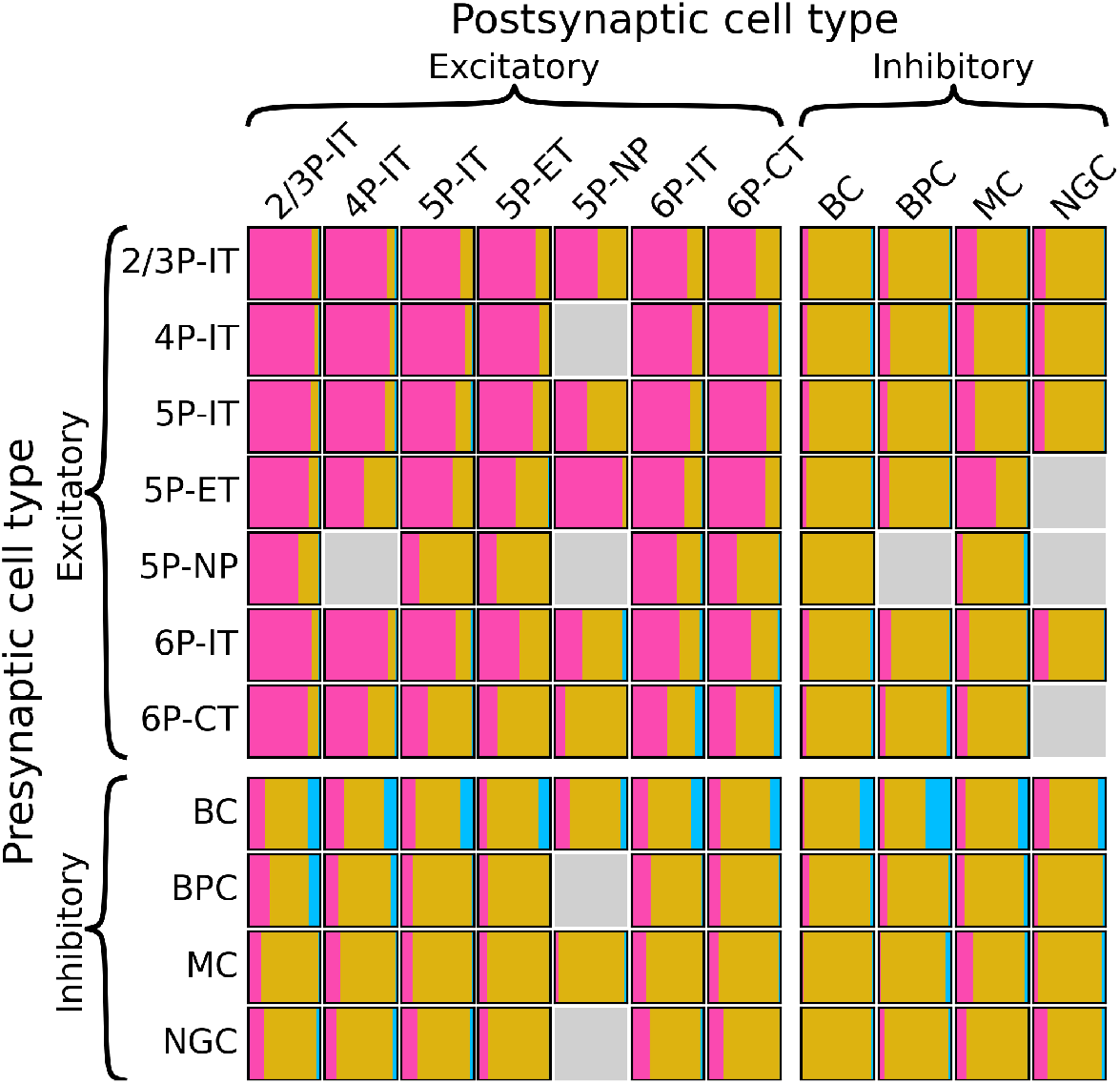
Description of within-column connectivity, showing the proportion of each cell type-to-cell type connection which connects to spine, shaft, or soma. Cell type connections with fewer than 10 synapses are shown as a grey box.

**Figure S6:**
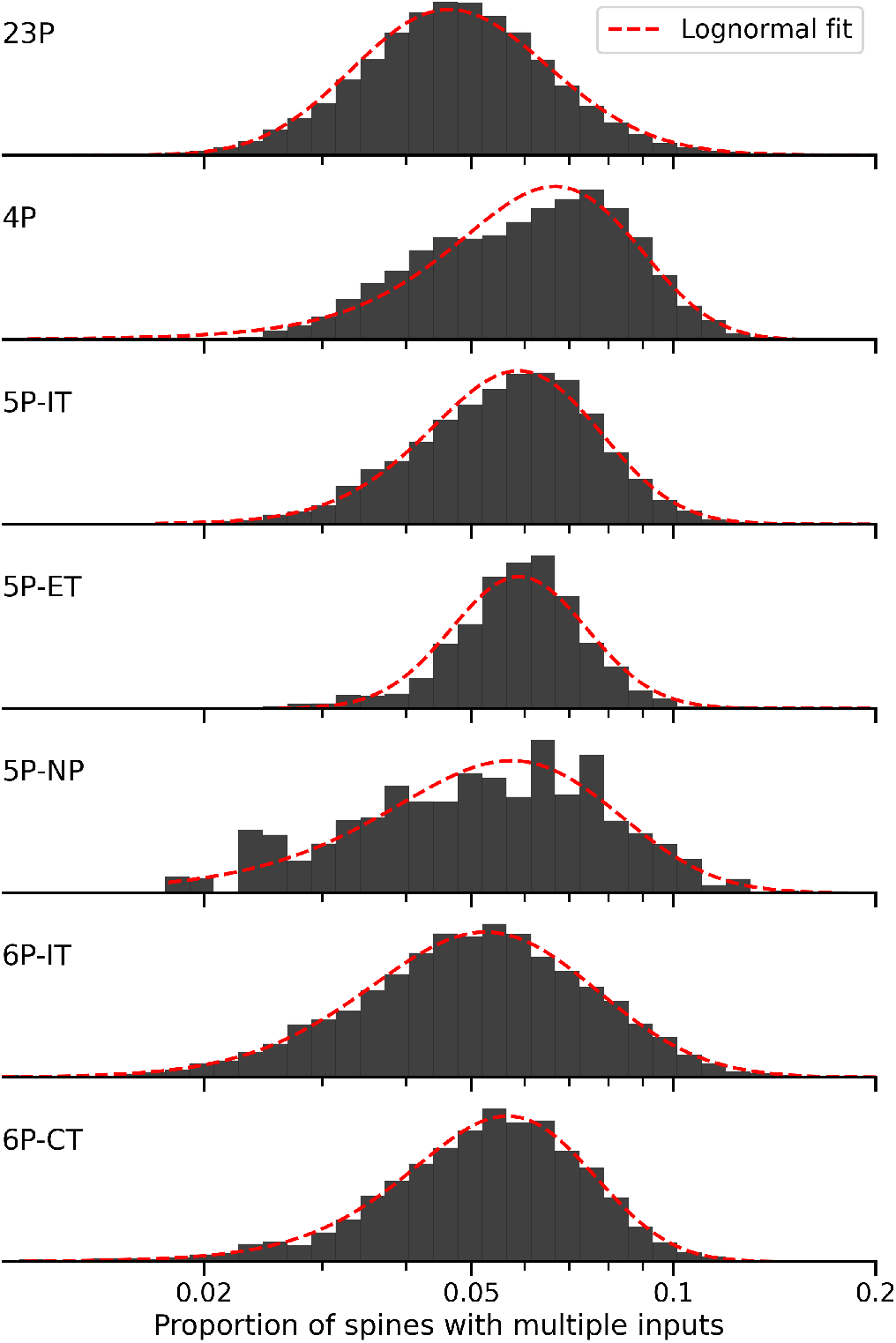
Distributions of proportion of spines with multiple inputs for excitatory neurons, stratified by major cell type. Red line shows best lognormal fit to each distribution.

**Figure S7:**
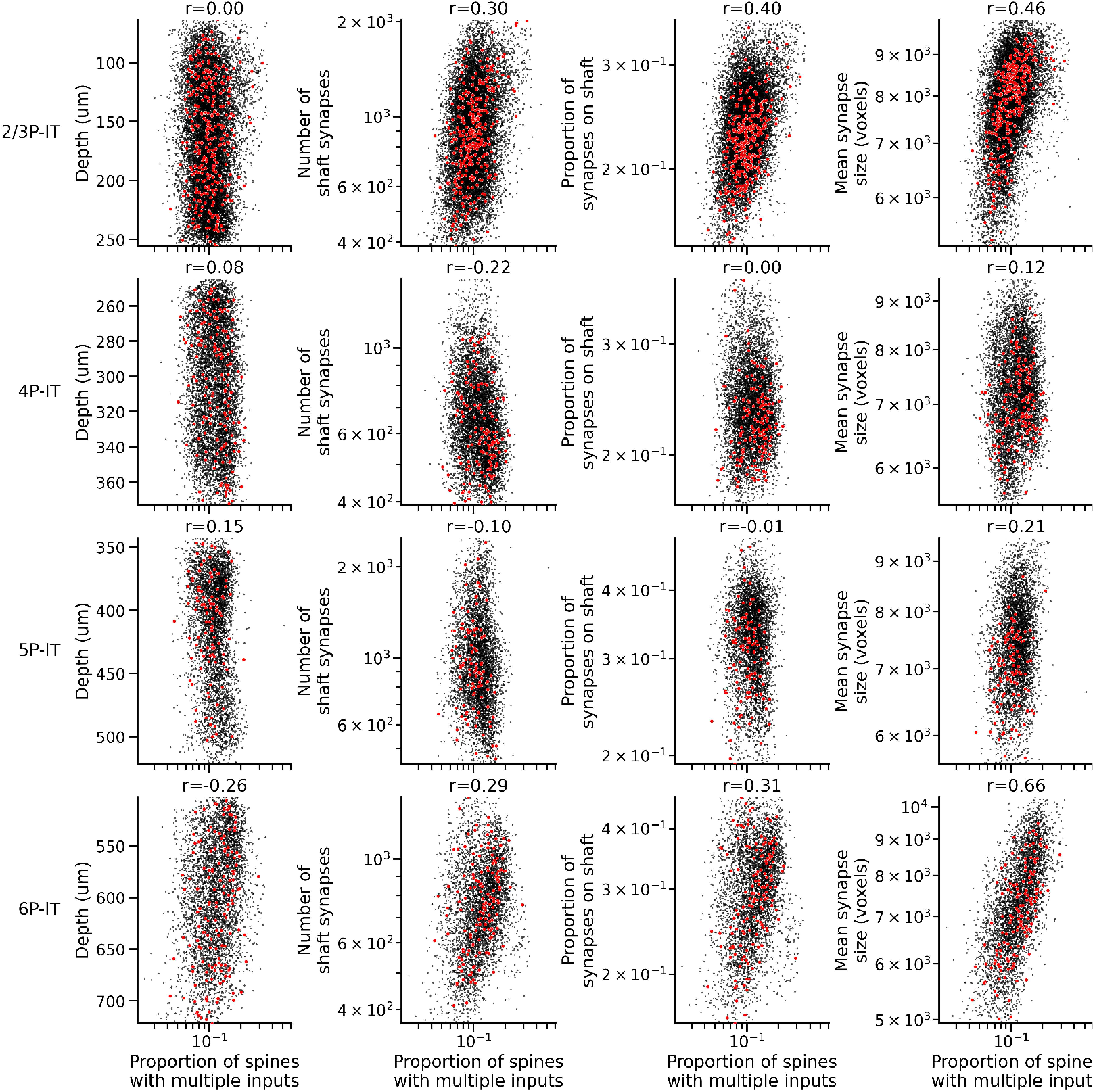
Relationships between a cell’s proportion of spines with multiple inputs (x-axis) with various other cell properties. Y-axes from left to right show: soma depth (lower number is closer to pia), total number of input synapses to shaft, proportion of input synapses to shaft, and the mean size of all synaptic inputs to that cell. Red dots indicate cells in the curated column. Panel titles show Pearson’s *r* correlation coefficient.

**Figure S8:**
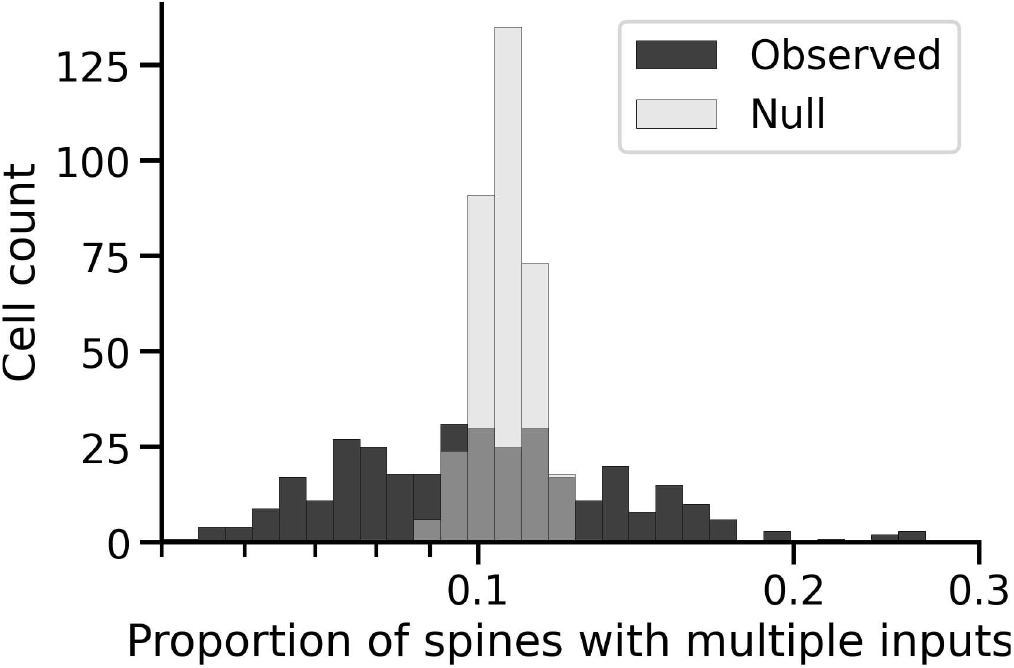
Distribution of proportion of spines with multiple inputs for Layer 2/3 pyramidal neurons (black) compared to a null model. The null model was constructed by first binning spine volume into 100 bins on a log scale, and then shuffling spine labels (as single- or multi-input) across cells within each bin. This procedure preserved the distribution of spine sizes per cell, but randomized which spines were multi-input. The grey curve shows the resulting distribution of multi-input spine proportions. Note that its cell-to-cell variance is much lower than observed, indicating that controlling for cell-to-cell differences in spine size alone does not explain the observed variability in multi-input spine rates.

**Figure S9:**
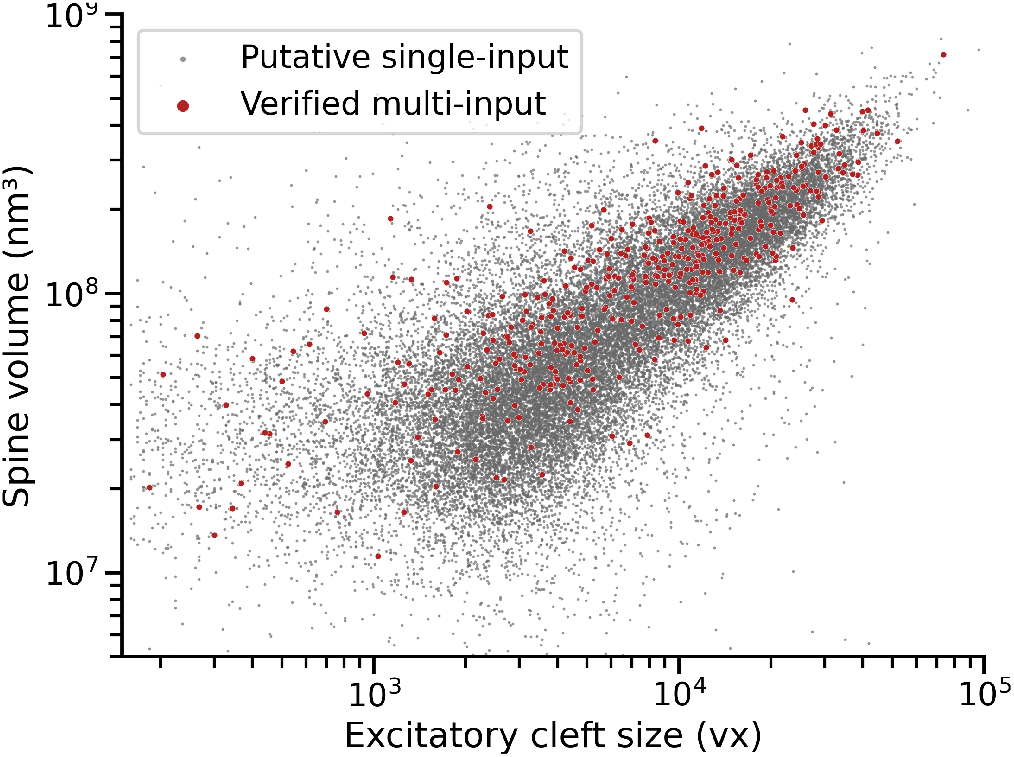
Correlation between the volume of a spine and the size of the excitatory presynaptic cleft onto that spine, as a function of whether the spine is single-input or multi-input. Multi-input spines are only shown in cases where we manually validated the unitary spine nature of that spine.

## Footnotes

1 github.com/bdpedigo/meshmash

2 github.com/bdpedigo/cloud-mesh

3 huggingface.co/bdpedigo/synapse_target_hks_rf

4 bdpedigo.github.io/neuron-hks/

5 bdpedigo.github.io/neuron-hks

## Notes

https://bdpedigo.github.io/neuron-hks/

### Competing Interest Statement

The authors have declared no competing interest.

### Summary of Updates

This version adds some information about running the proposed method on several other connectome datasets, including validation of model performance in one of these. It also includes various small changes to the text and wording.

### Funding

- National Institutes of Health: 5UM1NS132253-02

